# Contractile to extensile transitions and mechanical adaptability enabled by activity in cytoskeletal structures

**DOI:** 10.1101/2024.02.22.581411

**Authors:** Alexandra Lamtyugina, Deb Sankar Banerjee, Yuqing Qiu, Suriyanarayanan Vaikuntanathan

## Abstract

Cytoskeletal network architecture is crucial in determining emergent morphology and dynamics, particularly in processes like stress fiber nucleation, where cells respond to mechanical cues. However, a clear understanding of the connection between architecture, dynamics, and mechanical response remains lacking. In this study, we investigate how self-assembled cytoskeletal structures respond to external mechanical perturbations, focusing on filament and crosslinker mixtures in two dimensions. Using agent-based models complemented by coarse-grained thermodynamic analysis, we reveal how molecular motor activity enables cytoskeletal structures to robustly adapt to changing mechanical conditions. Our simulations demonstrate that under tensile forces, self-assembled active asters transform into bundle-like structures, reminiscent of de novo stress fiber formation in living cells, while pre-existing active bundles elongate further in a reproducible and regulated manner. In contrast, passive assemblies exhibit no such qualitative morphological reorganization, highlighting the critical role of activity in mechanical adaptation. We derive a simple relation approximating active stress as a function of average relative motor alignment, a measure of mesoscopic architecture, which allows us to predict changes in stress characteristics in response to mechanical perturbations with or without morphological reorganization. Our results demonstrate how mesoscopic architecture and motor activity work in tandem to enable robust morphological and mechanical adaptation in cytoskeletal structures, providing insight into cellular mechanosensing and stress fiber formation.

**SIGNIFICANCE:** Cytoskeletal networks adapt for optimal functionality in a dynamic intra-cellular environment. However, how adaptability emerges from local network architecture and their activity remains elusive. Here, we use specialized simulations and theory to study the response of cytoskeletal network structures, such as asters and bundles, to mechanical perturbation and show how activity enables robust morphological adaptation. Further, we show how the emergent contractile or extensile nature of these network structures is determined by a mesoscale architectural metric measured as average relative motor alignment. The mechanical adaptation emerges from changes in this architectural metric due to mechanical perturbations.

## INTRODUCTION

The cellular cytoskeleton self-assembles into various structures and networks that play crucial roles in the physiological functionality of the cell (1–4). Emergent forms and relative abundance of cytoskeletal structures are often determined by the mechanical environment of the cell (5–7), and correct structural organization of the cytoskeleton in response to its mechanical environment, such as substrate stiffness, myosin-generated pulsatile forces or external forces, is crucial for proper cellular functions (7–10). A striking example of mechanically-driven structural adaptation is stress fiber formation, where cells respond to mechanical cues–including myosin-generated contractile pulses and external forces–by nucleating and assembling contractile actomyosin bundles (11–15). Understanding how mechanical forces trigger such morphological transitions, and how molecular motors enable these adaptive responses, remains a fundamental challenge.

To address how self-assembled cytoskeletal structures respond to external mechanical perturbations, we examine the emergent dynamics of filament-molecular motor/crosslinker assemblies upon applying forces at the microscopic scale. Our agent-based simulations (Fig. 1A), complemented by non-equilibrium thermodynamic analysis, reveal how molecular motor activity allows cytoskeletal structures to respond robustly by changing their morphology to adapt to external forcing conditions. Inspired by experimental approaches that exert precise forces on cytoskeletal structures through optical activation of molecular motors (16) or external mechanical perturbations (17, 18), we develop a microscopic forcing protocol in which forces act on individual filaments along their long axis. Such forcing scenarios may also arise *in vivo* during tissue morphogenesis in which extrinsic forces act on cells and reorganize cytoskeletal networks (19, 20).

**Figure 1:**
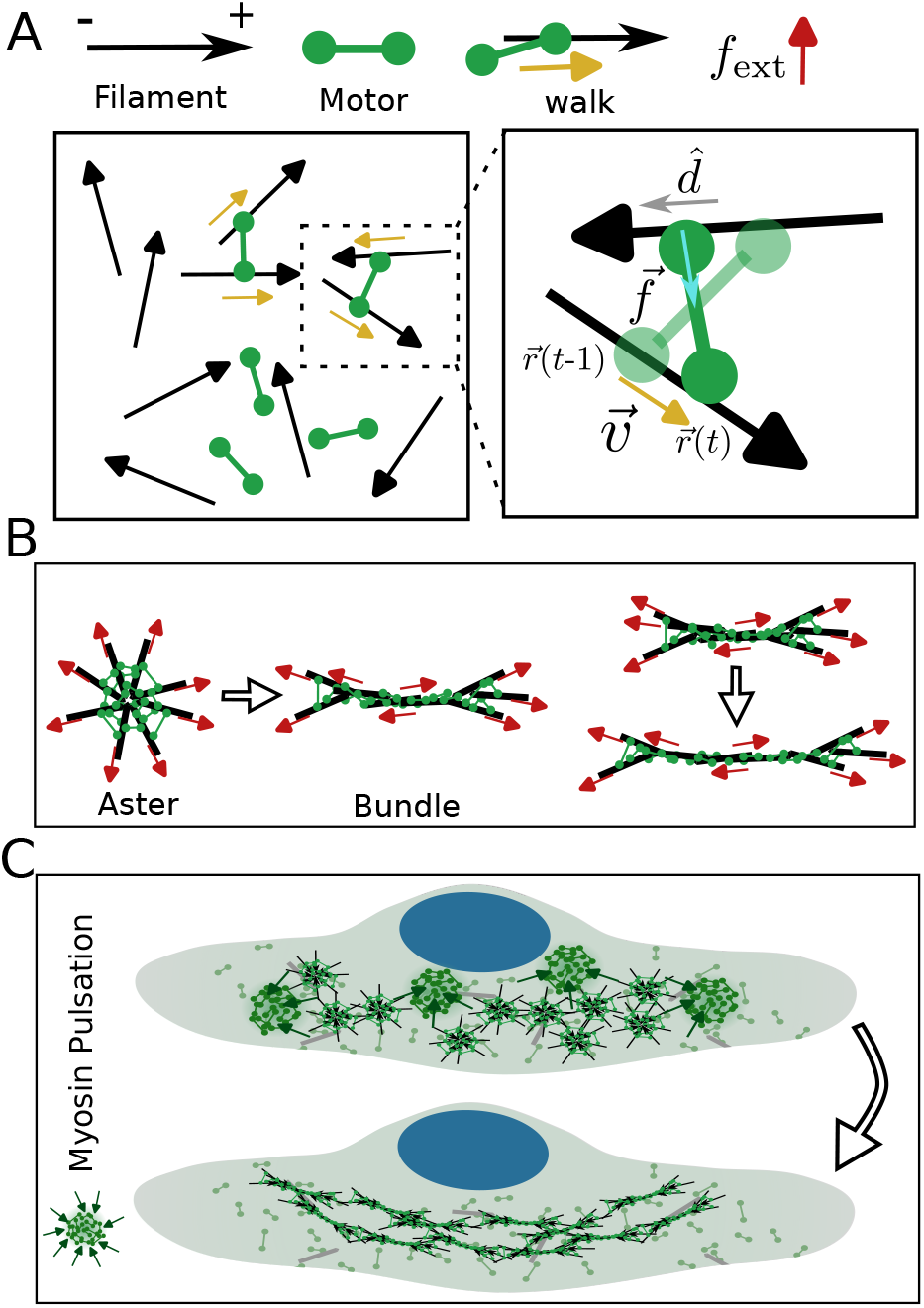
Self-assembly and response to mechanical perturbation of active cytoskeletal self-assemblies. (A) Schematic of polar filaments and motors that can bind and unbind from the filaments. The motors walk towards the plus or barbed end of the filaments (marked by arrowhead) with a speed v. (B) Schematic showing how asters and bundles respond to the externally applied force (*f*_ext_) and undergo morphological transformation and deformation, respectively. (C) Schematic of de novo cortical stress fiber formation in a cell in response to internally generated myosin pulsation (15), transient contractile events produced by myosin mini-filament clusters engaging the actin cortex (21), which generate local tensile forces that drive reorganization of the actin network into aligned bundle-like stress fibers.

Our simulations reveal qualitatively different mechanical responses between active (molecular motor-driven) and passive self-assemblies. Active structures resembling asters respond and adapt to external forces by transforming into bundle-like assemblies (Fig. 1B)—a transition reminiscent of de novo stress fiber formation in cells in response to forces (15) (Fig. 1C). Active bundle assemblies respond to external forces by elongating further (Fig. 1B). Under similar conditions, passive assemblies fail to mount comparably robust responses.

Our analysis reveals how these morphological transitions are accompanied by changes in the nature of active stress. We derive a simple scaling relation connecting the sign of active stress (contractile or extensile) to the mesoscopic architecture of the self-assembly. Notably, while typical aster-like active assemblies are contractile and typical bundle-like active assemblies are extensile, the bundles formed by force-induced transformation of asters retain contractile character and regain their original morphology upon force withdrawal. This innate response allows deformed active bundles to shrink back when external forces are removed. Together, our work provides a mechanistic framework for the force response and adaptation of cytoskeletal assemblies and suggests how pulsatile motor activity combined with mechanical tension may drive stress fiber nucleation.

## METHODS

We perform agent-based simulations of passive and active cytoskeletal assemblies with the Langevin dynamics-based Cytosim simulation engine (22). We examine a 2D system of filaments and crosslinking elements with various degrees of activity and apply an external force protocol inspired by the experimental setup of stretching an adhered cell by deforming the substrate. We ignore volume exclusion interactions to model a thin layer of filaments at low filament density, where their dynamics will depend on the network connectivity (23). Inspired by cortical actin filaments, the filaments in our simulation are modeled as short rigid rods (24, 25). While the cortical filaments show an exponential length distribution (26), for simplicity, we consider monodisperse filaments here. We ignore filament bending as the filament size is much smaller than their persistence length, leading to a high bending energy and negligible bending. We also ignore any bending of the cross-linkers and, assuming small elastic deformations for cross-linkers, we model them as two beads connected by a harmonic spring. The cross-linkers binding unbindings are modeled with a mechanosensitive slip-bond behavior. Active cross-linkers preferentially move towards one of the ends (the plus-end or barbed-end) of the polar filaments with a force-dependent speed 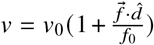 where 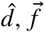, and *f*_0_ are a unit vector in the direction of filament plus-end/barbed-end, force in the motor, and the magnitude of stall-force, respectively (Fig. 1A). The concentrations of motors/crosslinkers and filaments are held constant. In each simulation trajectory, the largest connected cluster, defined as the set of filaments sharing at least one motor or crosslinker, was identified by filament count, and all structural analyses were performed on this cluster. For more details on the numerical simulations and relevant simulation parameters, refer to Section S.I in the Supplementary Information.

We modify Cytosim to be able to introduce arbitrary mechanical perturbations to the dynamics of system components. The mechanical perturbations take the form of external force vectors applied to individual filaments at their centers. The direction of the applied external force is determined by the polar filament’s orientation vector and its location relative to the whole assembly’s center of mass. The external force vector *f*_ext_ is chosen such that it is parallel to the direction of the filament and pointing away from the structure’s center of mass (Fig. 1B). The forcing protocol is designed to capture the mesoscale effect of substrate stretching on a self-assembled cytoskeletal structure. Under uniform substrate stretching, the net mechanical effect on an assembly is a radially outward tension, whose projection onto individual filament degrees of freedom yields outward forces along filament orientations. Since filaments are modeled as short rigid rods (*l* _*f*_ = 0.25*μ*m, much smaller than the assembly size), the distinction between force application at the filament center versus ends will be negligible. We therefore do not expect the qualitative results to depend on the precise force application point along the filament. We do not explore other forcing geometries in this work, but expect that radially outward forces applied at filament ends or boundaries would produce qualitatively similar morphological responses. Section S.II in the SI contains additional information on the implementation of the applied external forces.

### MORPHOLOGICAL TRANSITIONS IN PASSIVE AND ACTIVE CYTOSKELETAL STRUCTURES

To understand the role of activity in the morphological adaptation of cytoskeletal structures in response to the applied force, we first systematically explore how the dynamics and morphology of the filament and crosslinker self-assemblies evolve as we increase *activity* in the system. We study this numerically using the agent-based simulation discussed in the Methods section. Here, a crosslinker is modeled by setting the unloaded walking speed to zero (*v*_0_ = 0) in the expression of the force-dependent speed and the emerging self-assemblies at *v*_0_ = 0 are referred to as the passive filament-crosslinker self-assemblies.

Changing the concentration or mechanical properties of passive crosslinkers (or motors) can facilitate the formation of cytoskeletal structures like asters and bundles from an isotropic mixture of filaments and crosslinkers and motors (27– 32) and may lead to a transition from one morphology to another (33, 34). With increasing motor stiffness and walking speed the filament-motor structures undergo morphological transition from being asters to bundles (Fig 2). While the morphologies look similar in both passive (*v*_0_ = 0) and active (*v*_0_ > 0) cases, they have distinct structural differences. At low values of crosslinker/motor stiffness, the filaments and passive crosslinkers self-assemble into *apolar asters* where the filament heads randomly point inwards or outwards to the aster center (Fig. 2 & Fig. 3A). In contrast, with increasing walking speed, filaments and motors (i.e., active crosslinkers) self-assemble into active asters which are polarity-sorted structures with all filament heads pointing inwards to the aster center (Fig. 2 & Fig. 3C).

**Figure 2:**
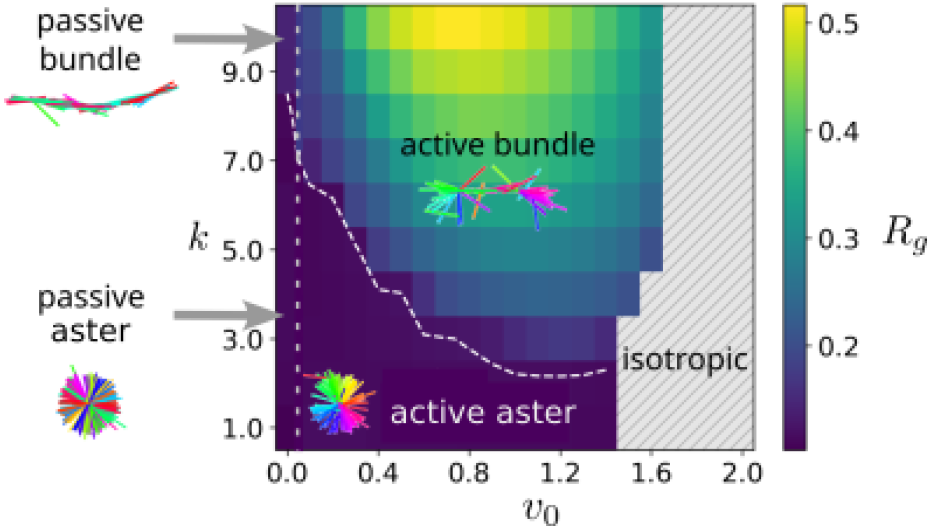
Structure regimes for varying unloaded walking speeds (*v*_0_) and crosslinker rigidities (*k*). In passive systems (*v*_0_ = 0), the structures do not exhibit polarity sorting. Adding activity to the cross-linking elements (motors) by increasing *v*_0_ results in similar structures but with additional order to filament orientations. Snapshots are color-coded to demonstrate filament orientation. The dashed white line denotes the boundary between aster and bundle regimes. The gray dotted vertical line denotes the boundary between passive (*v*_0_ = 0) and active (*v*_0_ > 0) systems.

**Figure 3:**
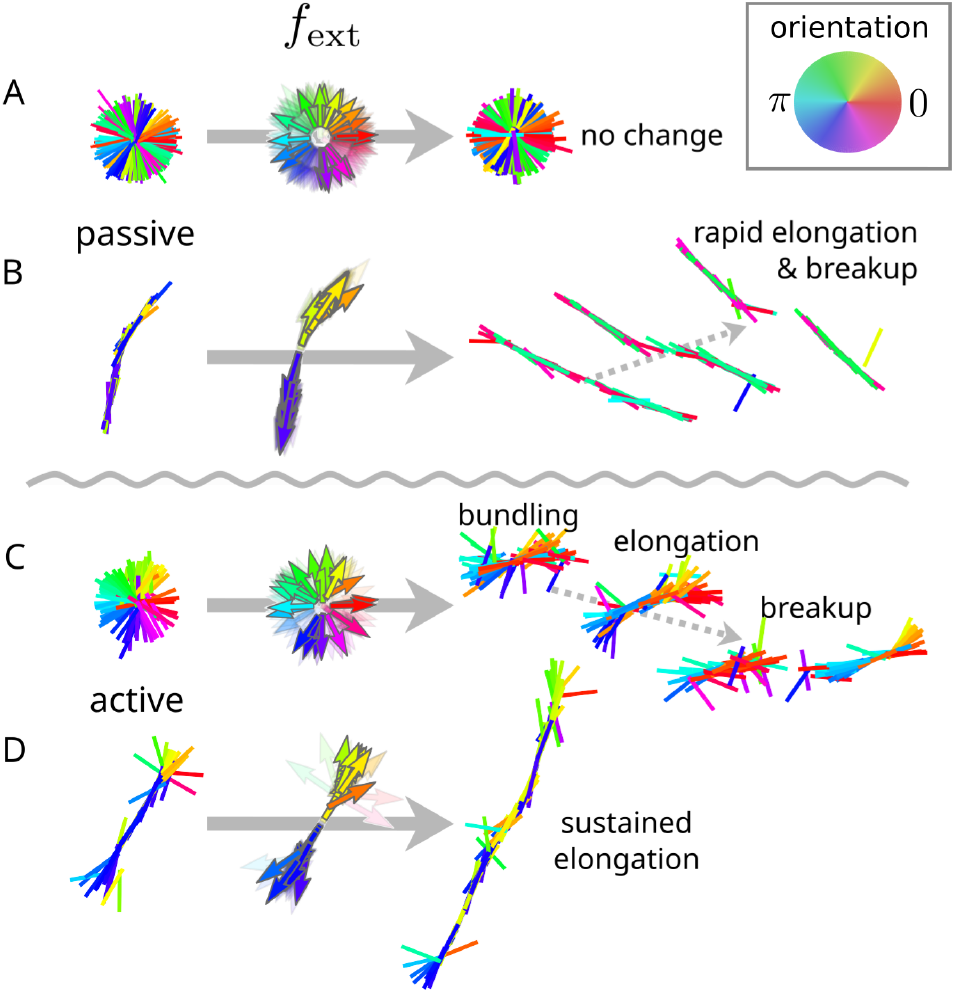
Passive (A, B) and active (C, D) structures respond differently to applied forces. Structures are color-coded based on filament orientations. External forces *f*_ext_ (arrows in center, also color-coded by orientation) are applied to the centers of each filament along filament orientation but always pointing away from the assembly’s center of mass. Dashed arrows indicate structure evolution as external forces continue to be applied.

The apolar asters transform into passive bundles with increasing crosslinker (i.e., *v*_0_ = 0) stiffness (Fig. 2). Passive bundles are structurally different from their active counterparts (at *v*_0_ > 0) with half-aster structures at the bundle ends and anti-parallel filaments more concentrated at the middle of the bundle rather than being distributed randomly along bundle length as in the passive case (Fig. 2 & Fig. 3B, D). In active bundles, the motors preferentially walk toward the barbed ends of filaments, driving the accumulation of anti-parallel filament overlaps at the bundle center via polarity sorting. In passive bundles, crosslinkers bind without directional preference, resulting in a uniform distribution of anti-parallel filament pairs along the bundle length. These underlying structural differences in passive and active cytoskeletal structures give rise to distinctly different responses to mechanical perturbations as we shall study in the next section.

### ACTIVITY LEADS TO ADAPTABILITY IN FILAMENT CROSSLINKER SELF-ASSEMBLIES

To study the response of the cytoskeletal structures to external mechanical perturbations, we apply an external force (*f*_ext_) on each filament of the self-assembled structures (see Fig. 3) and analyze the resulting morphological changes. We model the applied force pattern inspired by the forces experienced by a cell adhered to a substrate (please refer to the Methods and SI Section S.II for details of the force application protocol). Thus, at the mesoscopic level, the forces act to stretch the structures radially outward in the case of asters or axially outward in the case of bundles (Fig. 3). When perturbed with forces of similar magnitudes, the passive structures hardly show any qualitative change (in the case of passive asters, Fig. 3A) or are prone to fragmentation (in the case of passive bundles, Fig. 3B) while the active structures show much larger morphological changes and adapt to the new mechanical environment to maintain the structural integrity of the self-assembly (Fig. 3C, D).

We study the morphological transitions quantitatively by calculating the radius of gyration (*R*_*g*_) of the self-assemblies (the largest connected cluster was chosen in each case). The passive asters are the least sensitive to external forces and exhibit minimal changes to their radius of gyration (*R*_*g*_) (Fig. 4A) and remain in unaltered morphology (Fig. S1A).

**Figure 4:**
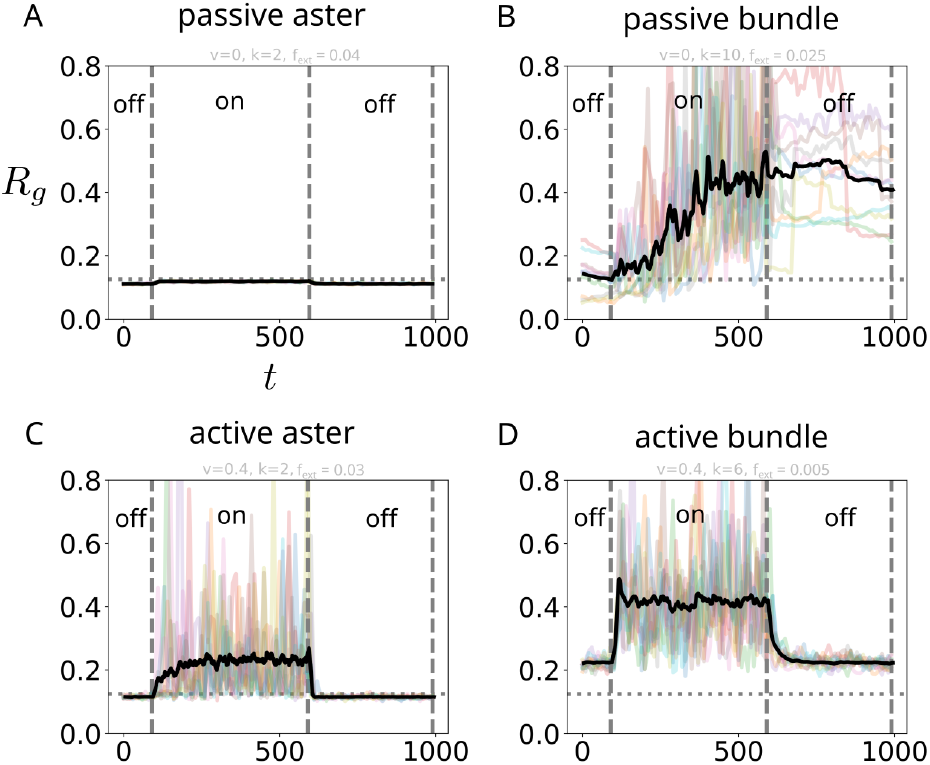
Traces of the radius of gyration (*R*_*g*_) of passive and active structures when a force is applied and maintained, and then released. Passive asters (A) are least responsive to external forces. Passive bundles (B) respond to forces, but do not maintain a steady state. Active asters (C) and bundles (D) achieve and maintain a new steady state upon the application of external forces. “On/off” labels indicate whether or not external forces are being applied at a given time regime (separated by vertical dashed lines). Horizontal dotted line indicates *R*_*g*_ value of aster configuration. Transparent color lines correspond to *R*_*g*_ traces for individual trajectories. Opaque black line is a running average across all trajectories in the ensemble.

The passive bundles on the other hand, are typically unstable under applied force. Passive crosslinkers binding and unbinding generate effective inter-filament friction (35), making passive bundles behave as viscous objects under sustained forcing. In the absence of an active restoring stress, the bundle deforms continuously until filaments are displaced beyond the crosslinker binding range, at which point fragmentation becomes irreversible. This viscous creep and rupture scenario is consistent with the rapid unsustained elongation observed across all force magnitudes in Figs. 4B and Fig. S1B–C In stark contrast, the active structures show reproducible morphological changes in response to the applied external force leading to the possibility of activity-driven mechanical adaptation of cytoskeletal structures. Active asters change morphology and form a *force-induced active bundle* under radially outward applied forces (Figs. 3, 4C, S2A, and S3A) and active bundles elongate to become larger bundles (i.e., with larger *R*_*g*_) in response to the axially outward applied forces (Figs. 3, 4D, S2B–C, and S3B). It is interesting to note that the active aster and active bundle both show recovery to their original morphologies once the applied force is removed (Fig. 4C & D). We revisit this apparently *elastic* behavior of active structures in a later section.

The force-dependent morphological transition from aster to the bundle is initiated above a critical threshold force. The precise value of this threshold force depends on different filament and motor (active crosslinker) parameters (Fig. S13). The computationally obtained *phase-diagram* of morphologies in motor stiffness (*k*) and external force magnitude (*f*_ext_) shows the critical force (marked by the boundary between aster and bundle phases) to decrease with increasing motor stiffness (Fig. 5A). This tunable force-dependent morphological transition signifies a potential mechanism for cellular mechanotransduction which may lead to stress fiber nucleation in response to mechanical cues. The active bundle elongation due to applied force also exhibits a regime where the elongation increases with applied force magnitude, but with increasing force, the bundle size generally decreases (Fig. 5A & B). This non-monotonic trend of *R*_*g*_ value saturation and decrease with increasing *f*_ext_ emerges from the crosslinker binding-unbinding kinetics, which is force-dependent and modeled as a slip-bond here. At higher values of external forces, the self-assembly starts losing filaments (i.e., the number of filaments, *N*_fil_ decreases) for all values of *k* due to increased motor disassembly and that leads to reducing *R*_*g*_ even though the morphology remains the same (Fig. 5C). It is important to note that the force-induced bundles (from active asters) described here represent a quasi-stationary state that is maintained as long as the external force is applied. Under sustained forcing, these bundles eventually may lose enough filaments over longer timescales due to force-accelerated motor unbinding (slip-bond kinetics), making them transient rather than permanently stable structures (Figs. S2–S3). This is distinct from mature stress fibers in living cells, which are actively maintained under mechanical load through continuous actin polymerization, myosin recruitment, and regulated crosslinker turnover, processes that are not included in our minimal model. Our results thus capture the initiation of a bundle-like state driven by tension, analogous to the early stages of de novo stress fiber nucleation (15), rather than the long-term maintenance of a mature stress fiber.

**Figure 5:**
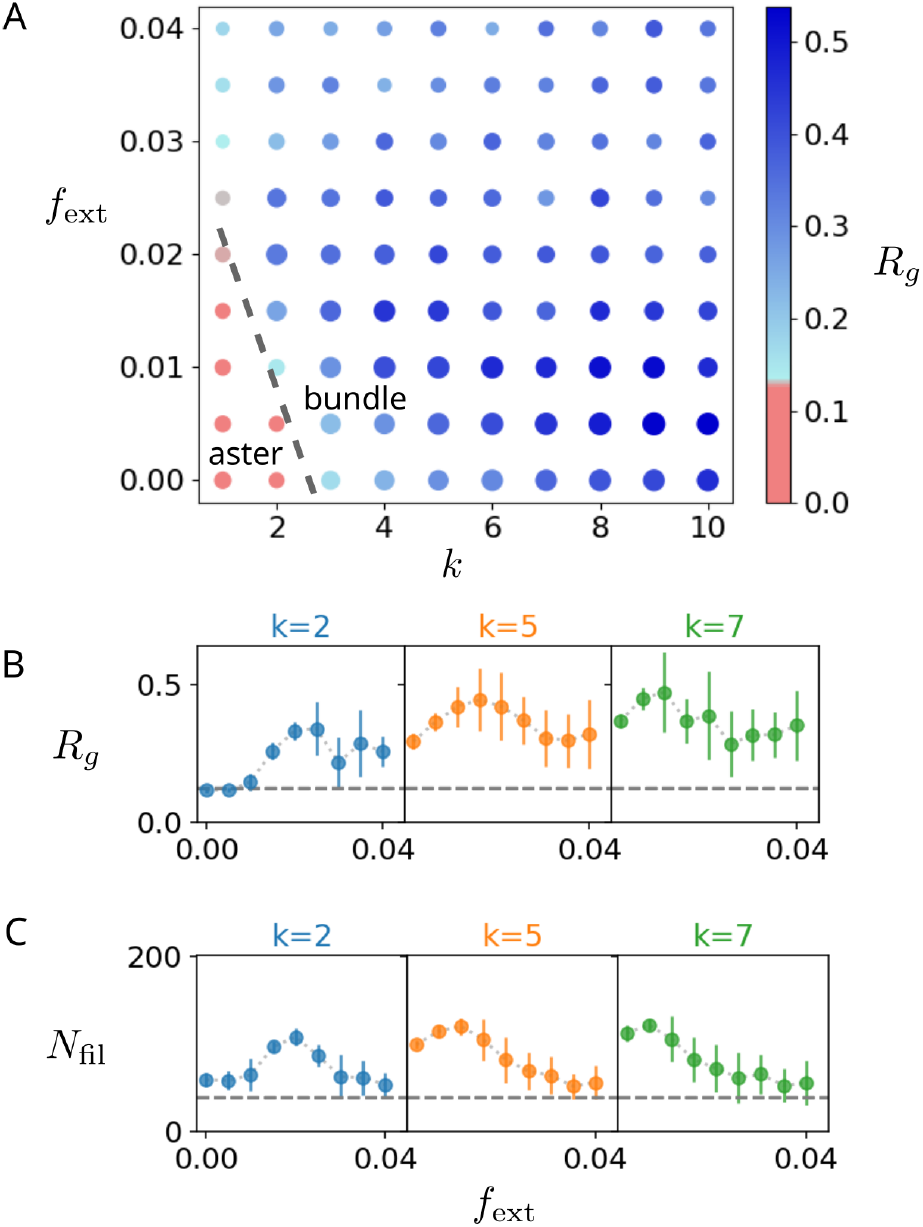
(A) Phase diagram upon the application of external forces (*v*_0_ = 1.2). Point color corresponds to *R*_*g*_, point size corresponds to cluster size, defined as the number of filaments in the cluster (*N*_fil_) normalized to the total number of filaments. (B) *R*_*g*_ for three values of *k* as a function of external force (*f*_ext_). Small values of *f*_ext_ result in structure elongation for both aster (*k* = 2) and bundle (*k* = 5, 7) starting states. (C) Number of filaments in the largest cluster (*N*_fil_) as a function of *f*_ext_. Large values of *f*_ext_ result in structure breakup, giving rise to lower *R*_*g*_ values.

### ACTIVE STRESS DETERMINES THE MECHANICAL RESPONSE OF ACTIVE SELF-ASSEMBLIES

Motor-dependent activity is important for the formation of contractile and extensile cytoskeletal structures (36–39). The results discussed in the previous sections describe how the non-equilibrium process of motor walking is also important for adaptability and mechanosensitivity in cytoskeletal self-assemblies. However, they do not provide an intuitive understanding of why the force response of the active self-assemblies resembles an elastic behaviour (i.e., force removal restores previous morphology). Here we argue that the answer lies in the active stress (of these mesoscopic structures) and its change in nature upon mechanical perturbation—changes that can be captured through a mesoscopic order parameter characterizing motor-filament organization. With motor walking being the only active process in our system, the active stress (*σ*^*a*^) generated in the self-assembled mesoscopic structures can be estimated from the average rate of work done or average power input as

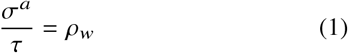

where *τ* is a characteristic timescale which we take to be the average motor residence time 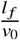 and *ρ*_*w*_ is the density of rate of work done which can be defined over a small area Ω_*A*_ as

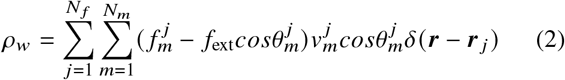

where the index *j* represents the filaments in the area Ω_*A*_ and *m* represents the index of motors bound to a filament. Here 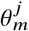 is the relative motor alignment, i.e., the angle between *m*^th^ motor axis and the polarity direction of *j* ^th^ filament. We arrive at the above relation by using the definition of the average rate of work done or average power input: 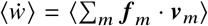 and the modified motor force in presence of applied force (*f*_ext_) given by 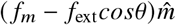, where 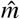 is the motor axis direction.

In general, the motor forces, motor alignment and walking speed are functions of the local filament configuration and network connectivity but for simplicity, we replace them with respective average quantities (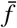, ⟨*cosθ*⟩ and 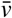) to arrive at

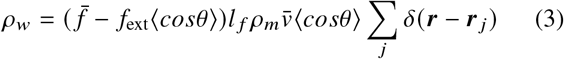

where we have replaced the number of motors attached to a filament (*N*_*m*_) with *l* _*f*_ *ρ*_*m*_, *ρ*_*m*_ being the average motor density on filaments. We apply this mean-field consideration assuming the structures are densely connected and the motor forces and motor velocities are narrowly distributed around their mean value such that the mean value capture the typical values of these quantities. Further breaking down the average motor force and walking speed in terms of system parameters as 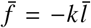 and 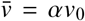 (although *α* is a function of system parameters and network configurations, we assume it to be constant) we arrive at a definition of active stress as

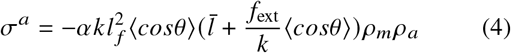

where *ρ*_*a*_ is the local filament density. In our derivation, the active stress is computed by summing over all filaments, yielding one factor of *ρ*_*a*_. The motor density on filaments, *ρ*_*m*_ is a fixed parameter, rather than being set by equilibration with a reservoir. If instead one considers the density of motors successfully bridging two filaments, this would scale as *ρ*_*a*_ *ρ*_*m*_, recovering previously reported 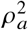 dependence (40). We can rewrite the active stress as *σ*^*a*^ = *ξ ρ*_*m*_*ρ*_*a*_ with *ξ* < 0 indicating contractile activity and *ξ* > 0 indicating extensile activity. Here the sign of the *contractility* parameter 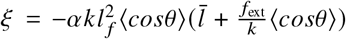 is determined by the sign of ⟨*cosθ*⟩ as all other quantities are positive and typical values of 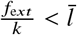. If we consider the geometry of motor connections in the aster, then we can argue that ⟨*cosθ*⟩_Aster_ > 0 as most of the motor connections (in both parallel and anti-parallel filament pairs) have *θ* < *π*/2 (Fig. 6A).

**Figure 6:**
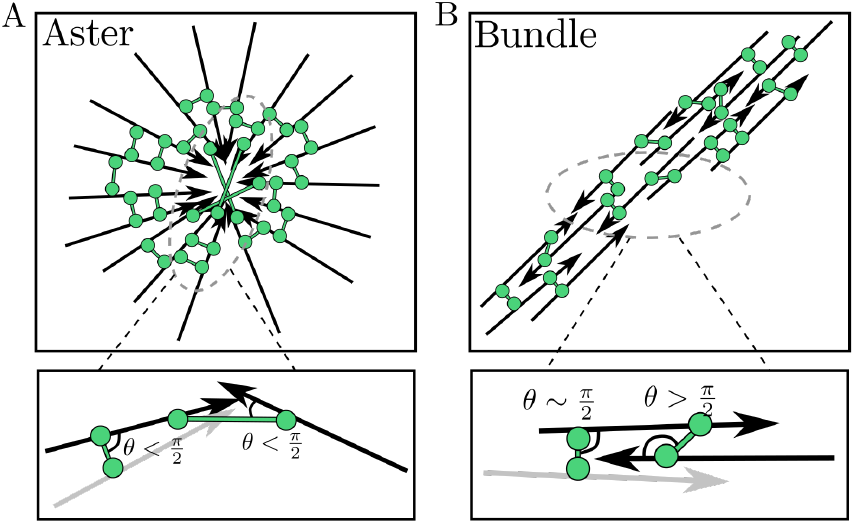
Filament and motor connections in aster and bundle. (A) Filament configuration and motor connectivity in an aster show angle between the motor axis and filament is smaller than *π*/2. (B) Filament configuration and motor connectivity in a bundle show angle between the motor axis and filament is larger than *π*/2.

This leads to *ξ* < 0 indicating the active stress for asters being contractile in nature. Conversely, in bundles, motor connections mostly have *θ* > *π*/2 in anti-parallel filaments while motor connections in parallel filaments do not contribute significantly as *θ* ∼ *π*/2 (Fig. 6B). This leads to ⟨*cosθ*⟩ < 0 giving rise to extensile active stress (*contractility* parameter *ξ* > 0) in active bundles. Quantification of the ⟨*cosθ*⟩ from simulations shows a clear change in sign as the active aster changes into a bundle with increasing motor stiffness (Fig. 7A). This indicates a change in the nature of active stress, from being contractile in aster to being extensile in bundles, associated with the morphological change. The architectural quantity, ⟨*cosθ*⟩ characterizing the relative motor-filament alignment, acts as a mesoscopic order parameter for the contractile-to-extensile transition in active stress. These active extensile bundles reach a dynamic steady state through filament binding-unbinding. The bundles have both parallel and anti-parallel filament pairs connected by motors. Anti-parallel motor pairs drive filament sliding, progressively expelling filaments from the overlap zone. Expelled filaments encounter neighboring filaments and form new parallel or anti-parallel pairs, while filaments at the bundle boundary unbind when they slide and are continuously recruited back into the bundle through motor binding. A stationary mean bundle size emerges when the rate of filament expulsion from the anti-parallel sliding balances the rate of filament recruitment (or rebinding). This is a dynamic steady state in which sliding continues but is sustained by continuous filament binding and unbinding, conceptually consistent with the flux balance framework of Kruse & Julicher (40) (see Section S.VII in the SI for additional details).

**Figure 7:**
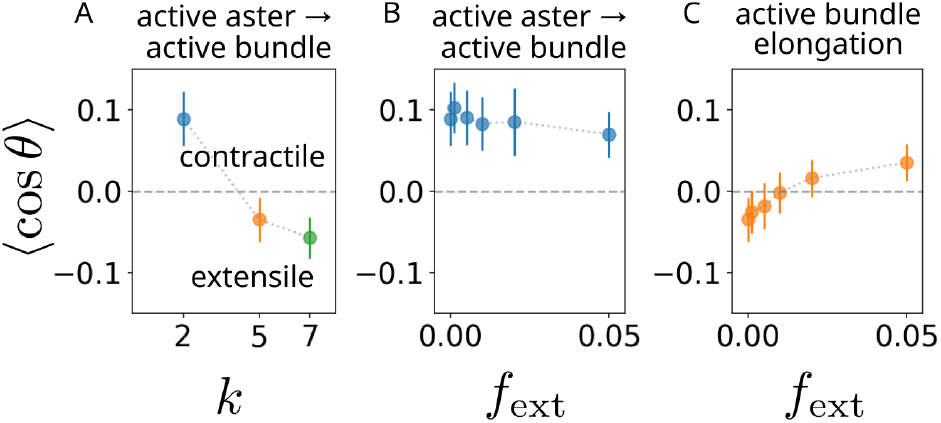
⟨cos *θ*⟩ serves as a proxy for quantifying the mesoscopic architecture of filament-crosslinker assemblies (where *θ* is the angle between a motor and a filament). For ⟨cos *θ*⟩ > 0, structures exhibit contractile behavior, whereas for ⟨cos *θ*⟩ structures exhibit extensile behavior. Point colors correspond to different *k* (blue: *k* = 2, orange: *k* = 5, green: *k* = 7). (A) The aster to bundle transition achieved through changing motor rigidity *k* results in change in sign of ⟨cos *θ*⟩ . (B) The morphological transition from aster (*k* = 2) to bundle as a result of applied *f*_ext_ is not accompanied by a contractile-extensile transition, resulting in the bundle contracting back to aster form upon the removal of *f*_ext_. (C) The elongation of initially extensile active bundles (*k* = 5) results in contractile structures that return to their original state when external forces are removed.

Further, the change in ⟨*cosθ*⟩ due to mechanical perturbation displays qualitative differences in active asters and active bundles. In contractile active asters, the external forces do not change the sign of ⟨*cosθ*⟩ , indicating that the bundles obtained by applied forces are contractile in nature (Fig. 7B). Due to this contractile nature, the bundles created from deforming asters shrink back and reorganize into asters when applied forces are removed, leading to restoration of the original morphology (Fig. 4C). The applied forces lead to a change in sign of ⟨*cosθ*⟩ from negative to positive in active bundles (Fig. 7C), indicating the bundles become contractile when external forces are applied. This change in the nature of active stress explains the restoration of bundles to their original length when applied forces are removed (Figs. 4D, S2, and S3). The timescale for this self-regulation is set by the motor traversal time *τ*_rev_ ∼ *l* _*f*/_*v*_0_ ∼ 0.2 *s* (Supplementary Table I), which is fast relative to the forcing window (500 *s*), explaining the rapid attainment of a new steady-state *R*_*g*_ in Fig. 4D.

The changes in the ⟨*cosθ*⟩ in response to applied forces emerge from collective dynamics and depend on the details of the architecture of the filament-motor self-assemblies. Although previous theoretical studies have predicted extensile or contractile behaviour based on network architecture (41–44), we believe our study is the first to report a force-dependent transition between extensile and contractile structures. Finally, while a full theoretical treatment of these changes is challenging, we present a simple set of arguments in an attempt to explain the observed changes. Due to the architecture of the asters, external forces do not change the sign of the *cosθ* but in bundles the external outward forces make the angles cross *π*/2 in motors between anti-parallel filaments, causing a change in the sign of *cosθ* (Fig. 8).

**Figure 8:**
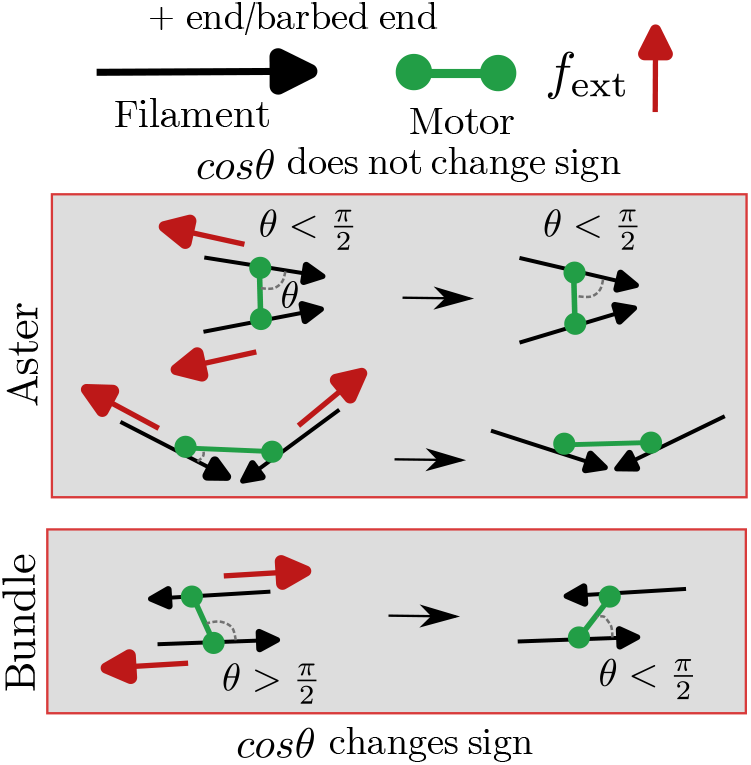
Force-dependent changes in self-assembly architecture. Forces acting on filaments in aster (top) and bundle (bottom) show the change in the relative motor alignment (*θ*) for two-filament example leads to a change in the sign of *cosθ* for bundles but not for aster.

## DISCUSSION

In this work, we study how cytoskeletal structures respond to mechanical perturbations and the role of activity in eliciting such responses. Recent experiments have shown that cells can nucleate stress fibers de novo from actomyosin condensates through pulsatile myosin activity and mechanical tension (15), yet the physical principles underlying such mechanically-driven morphological transitions remain unclear. Although we consider a simplified description of the cytoskeleton composed of small rigid filaments and (active or passive) crosslinkers, our simulations reproduce the emergence of asters and bundles observed in previous computational and experimental studies (28, 45–47).

We model external forces acting radially outward (or axially for bundles) to mimic forces on the cytoskeletal network of adhered cells on stretched substrates, and find qualitatively different responses between active and passive structures. Active asters transform into bundles above a threshold force—reminiscent of stress fiber nucleation—while active bundles elongate in a regular manner. In contrast, passive asters show no qualitative change and passive bundles disintegrate. This programmable, switch-like response demonstrates how motor activity enables mechanosensation and structural adaptation.

A key finding is that morphological transitions are accompanied by changes in active stress. Bundles formed by force-induced transformation of asters are contractile and restore their original morphology upon force removal—unlike the extensile bundles formed by increasing motor stiffness. This force-induced contractility provides mechanistic insight into how cells nucleate and maintain contractile stress fibers under mechanical load, with motor-induced activity giving rise to elastic behavior at the mesoscopic scale (48).

Extensile to contractile transition in active stress has been recently found in experiments with actin-myosin systems (49) and with microtubule-kinesin systems (50–52) but the physical mechanism remains poorly understood. In our case, we find contractile to extensile transition in active stress (associated with a morphological transition from aster to bundle) with increasing motor walking speed (Fig. 2). This is consistent with contractile to extensile transition observed in the microtubule-kinesin system with increasing ATP (chemical fuel for motor walking) (51, 52) . While competition between nematic alignment and polar sorting has been proposed as a controlling mechanism for this transition (51), our results indicate force-dependent architectural changes can also trigger this transition without any explicit way of affecting nematic alignment (as no volume exclusion between filaments is considered in our system). Notably, we observe extensile-to-contractile transition during bundle elongation without overall morphological change, demonstrating that active stress transitions are not always coupled to morphological reorganization.

Throughout this work, ⟨*cosθ*⟩ has been defined and computed in two dimensions, where the angular distribution of motor orientations relative to filament polarity has a simple planar structure. This quantity generalizes naturally to three dimensions as *θ* remains the angle between the motor axis and the polarity direction of the filament to which it is bound, and ⟨*cosθ*⟩ is averaged over all bound motors in a volume element. If the active stress is treated as isotropic, then the scalar ⟨*cosθ*⟩ remains a complete descriptor of the contractile or extensile character of the assembly in three dimensions, with the same sign convention (⟨*cosθ*⟩ > 0 contractile, ⟨*cosθ*⟩ < 0 extensile) as derived here. The richer tensorial structure of active stress would only need to be accounted for in situations where stress anisotropy is significant.

In standard active gel theory, the active stress is written as *σ*_*a*_ = *ξQ*, where *ξ* is a constant activity coefficient whose sign distinguishes contractile (*ξ* < 0) from extensile (*ξ* > 0) systems. Comparing with our Eq. 4, the effective activity coefficient in our system is 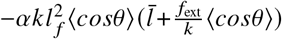, revealing two features absent from standard continuum descriptions: effective *ξ* depends on the mesoscopic motor-filament alignment ⟨*cosθ*⟩ , which evolves with assembly architecture, and it carries an explicit dependence on the external force *f*_ext_ that can switch its sign. To capture such force-induced contractile-to-extensile transitions, continuum theories would need to track a polar order parameter analogous to ⟨*cosθ*⟩ and couple its dynamics to both the nematic order field and the external stress — an extension that our microscopic results directly motivate.

Our results are qualitatively robust to two specific model variations tested: catch-bond crosslinker dynamics and filament length polydispersity. Simulations with catch-bond dynamics and filament length polydispersity (exponential distribution between 0.15 − 0.35 *μ*m compared to monodisperse 0.25 *μ*m filaments) show qualitatively similar morphological transitions (Fig. S14-17) to the baseline case with slip-bond behavior and monodisperse filaments (Fig.5). The emerging self-assembly structures are robust under variation in motor and crosslinker density as well. No stable structures form at low density of motor and crosslinkers and with increasing density, we find asters and bundle structures (see Supplementary section S. VIII and Fig. S18-19). This demonstrates the broad applicability of our findings.

Experimental data for a direct comparison with our findings are currently lacking but the model predictions can be tested in future experiments. The key predictions are: (i) A force-dependent, reversible aster-to-bundle morphological transition in active cytoskeletal self-assemblies. This transition occurs above a threshold applied force (dependent on motor stiffness and other system parameters), with active asters transforming into aligned bundle structures. Upon force removal, these bundles exhibit elastic recovery, shrinking back and reorganizing into their original aster morphology—a behavior absent in passive assemblies. (ii) The transition of active stress from extensile to contractile character in bundle structures subjected to external forces. While bundles formed by increasing motor stiffness are extensile, those formed by force-induced transformation of asters retain contractile character. This force-dependent stress reversal, mediated by changes in mesoscopic motor alignment, explains the elastic mechanical response and provides mechanistic insight into cellular stress fiber maintenance under load.

Possible experimental approaches include: attaching beads to reconstituted actomyosin clusters and applying constant outward force using multi-trap optical tweezers (53) to measure morphological changes; embedding actomyosin networks in soft gels or on elastic posts and stretching the substrate (54); or using magnetic beads to pull the network. High-resolution fluorescence imaging could quantify structural changes, while rheological measurements (e.g., active microrheology (55)) or laser ablation (56) could detect predicted shifts in rigidity or tension when active stress switches from extensile to contractile.

In summary, we demonstrate how self-assembled cytoskeletal structures undergo morphological transitions in response to mechanical perturbations, with activity playing a crucial role in determining mechanical response. The force-induced aster-to-bundle transition, coupled with the emergence of contractile character in the resulting bundles, provides a mechanistic framework for understanding tension-induced stress fiber nucleation (15). While our system is simplified, we anticipate broad applicability to cellular force sensing, stress fiber dynamics, and cytoskeletal remodeling during morphogenesis. An important direction for future theoretical work is developing hydrodynamic theories that can capture active stress transitions from extensile to contractile—a phenomenon not yet adequately described by existing frameworks.

## DATA AVAILABILITY

Simulation input files, Cytosim parameter files, and analysis scripts that support the findings of this study are available from the corresponding author upon reasonable request.

## AUTHOR CONTRIBUTIONS

A.L., Y.Q. and D.S.B. designed research with input from S.V. A.L. and Y.Q. performed simulations. A.L., D.S.B., and Y.Q. analyzed data. A.L. and D.S.B. developed the theoretical analysis with guidance from S.V. A.L., D.S.B., and S.V. wrote the manuscript.

## DECLARATION OF INTERESTS

The authors declare no competing interests.

## ACKNOWLEDGEMENTS

We thank Aaron R. Dinner for comments on the manuscript. A.L., D.S.B, Y.Q., and S.V. were supported by the National Institute of General Medical Sciences of the NIH under Award No. R35GM147400.

## Notes

### Competing Interest Statement

The authors have declared no competing interest.

### Summary of Updates

Result section updated to remove thermodynamic theory; Model section and Figure 1 updated to clarify forcing protocol; Figure 6 … Figure 9 removed as we removed thermodynamic model.

