## Supplementary Information for "Contractile to extensile transitions and mechanical adaptability enabled by activity in cytoskeletal structures"

Alexandra Lamtyugina*,[1] Deb Sankar Banerjee*,[1,2] Yuqing Qiu,[1,2] and Suriyanarayanan Vaikuntanathan†[1,2]
[1] *Department of Chemistry, University of Chicago, Chicago, IL 60637*
[2] *James Franck Institute, University of Chicago, Chicago, IL 60637*

### I. SIMULATION DETAILS

We carry out agent-based simulations of cytoskeletal structures using Cytosim [1]. In this system, motors are modeled as harmonic springs and filaments as short rigid rods. The motors exhibit slip-bond behavior, with a velocity that is determined by the force experienced by the motor head:

$$v = v_0 \left(1 + \frac{\vec{f} \cdot \hat{d}}{f_0}\right) \tag{1}$$

where $\hat{d}$, $\vec{f}$, and $f_0$ are a unit vector in the direction of filament plus-end/barbed-end, force in the motor, and the magnitude of stall-force, respectively. To change the degree of activity that the motors in the system exhibit, we modulate the $v_0$ parameter.

Each system was initialized to have filaments and crosslinkers/motors randomly distributed throughout the simulation space. The systems were then simulated for 2100 s without the application of external forces to allow for the formation of a steady-state structure to which the mechanical perturbations will be applied. Forces were then applied for 500 s and released, followed by 400 s of simulation time without applied forces to allow structures to respond to the removal of forces. The data sets in this work have an ensemble size of $N = 50$ for each tested parameter set.

Simulation parameters are enumerated in Table I.

For additional details on the Cytosim simulation engine, refer to [1].

### II. MODIFICATIONS TO CYTOSIM TO INTRODUCE EXTERNAL FORCE PERTURBATIONS

In order to introduce arbitrary mechanical perturbations to the system dynamics, we added a Python interface to Cytosim in order to directly manipulate the forces experienced by the filaments at each simulation step. The interface was implemented through the `ctypes` library [2]. The source code for the interface can be found on GitLab [3] in the `ctypes/` subdirectory of the repository.

The interface is written to allow for the application of arbitrary force vectors to arbitrary points on filaments. We specifically implement a protocol in which the force vectors are applied to the centers of filaments along filament orientation and pointing away from the structure's center of mass (see Fig. 2 in main text). A structure is defined by the filaments directly and indirectly connected to each other by cross-linkers/motors. We apply the force protocol only to the filaments belonging to the largest structure in the simulation, and all analyses are shown for the largest structures only.

### III. SNAPSHOTS AND MOVIES OF PASSIVE AND ACTIVE STRUCTURES RESPONDING TO APPLIED EXTERNAL FORCES

Representative snapshots of passive and active systems under different $f_{\text{ext}}$ conditions are shown in figures 1, 2, and 3.

---

* These authors contributed equally to this work.

In passive systems ($v_0 = 0$), aster starting states do not undergo any structural transformation upon the application of forces in the regimes examined by this study (Fig. 1A). Passive bundles (Fig. 1B,C) undergo elongation, but the elongation is not sustained even for small forces ($f_{\text{ext}} < 0.01$). Passive bundles continue to elongate until break up of the structure occurs. Since the systems contain a constant number of filaments and cross-linkers, beyond a certain degree of extension the number of cross-linkers per unit length of structure becomes insufficient to hold the structure together through passive cross-linking only.

The structures' response is different in the presence of active cross-linkers (Figs. 2 and 3). Active asters (Figs. 2A and 3A) undergo an initial elongation transformation into a bundled state. However, as the external forces continue to be applied, the newly-formed bundle state undergoes breakup and the smaller structures undergo further elongation. Once the forces are released, the system returns to its original polar aster steady state.

Active bundles (Figs. 2B,C and 3B) also undergo elongation. Active bundles are more sensitive to the magnitude of $f_{\text{ext}}$, undergoing elongation at much lower force magnitudes than active asters. The breaking of radial symmetry in active asters requires a higher magnitude of $f_{\text{ext}}$ than is required for the elongation of an already-formed bundle (which is why the bundle that forms after an aster is elongated in Figs. 2A and 3A eventually breaks up). For low force values, an active bundle is able to achieve a stable elongated steady state, as seen in Fig. 2B. When external forces are removed, the artificially elongated bundle contracts to its original length. For large $f_{\text{ext}}$, active bundles elongate but eventually break up (Figs. 2C and 3B). Although at a given instant the new elongated steady state is not stable for all values of $f_{\text{ext}}$, on average the structures under the application of forces are longer.

### IV. CHARACTERIZING STRUCTURES

All of the structure characterizations in this study were performed on the largest structure (i.e., the structure with the most number of filaments). Structure clustering is a built-in function of Cytosim and was used to determine the filaments belonging to the largest cluster.

#### A. Radius of gyration

To quantify the length of an aster or bundle structure, we employ the radius of gyration ($R_g$):

$$R_g = \sqrt{\frac{1}{N}\sum_{i=1}^{N} r_i^2} \tag{2}$$

where $r_i$ is distance between the center of mass of filament $i$ and the center of mass of the cluster to which filament $i$ belongs. $R_g$ is a convenient quantity to use without needing to calculate the absolute length of the structure, which becomes difficult for longer bundles which may curve at multiple points along their axial length. Fig. 4 summarizes the trends in $R_g$ for various levels of activity ($v_0$), motor rigidities ($k$), and external forces ($f_{\text{ext}}$).

#### B. Number of filaments and motors

The number of filaments $N_f$ (Fig. 5) is useful as it allows us to quantify the integrity of the structures as they respond to external forces. In both passive and active cases, structures begin to disintegrate for large enough force magnitudes, which is captured by a decrease in $N_f$.

Figure 6 shows the average number of motors in the largest cluster under various motor activity and rigidity conditions.

#### C. Motor-filament angles

By quantifying the change in orientation of active cross-linkers with respect to the filaments to which they are bound we can capture the change in the mesoscopic architecture of structures as they adapt to external forces (Fig. 7).

#### D. Motor extension

The length of cross-linkers provides insight into the forces that they experience as the structures adapt to different forcing conditions. In active bundles, average motor extension increases with $f_{\text{ext}}$ (Fig. 8).

### V. FORCE EXTENSION DIAGRAMS FOR VARIOUS LEVELS OF MOTOR ACTIVITY

Figure 9 summarizes the response of passive and active structures to various degrees of external forcing. Passive cross-linkers are not very responsive to external forces, especially for asters (low $k$m Fig. 9A).Introducing activity to the cross-linkers results in more responsive structures. Asters elongate into bundles at high $f_{\text{ext}}$, and bundles elongate at lower values of $f_{\text{ext}}$ (Fig. 9B–D). Bundles undergo structural breakup, as evidenced by smaller point sizes for increasing forces.

### VI. TIME-EXTENSION PLOTS OF $R_g$ FOR SYSTEMS UNDERGOING MECHANICAL PERTURBATIONS

#### A. Passive structures

Passive asters do not experience an increase in $R_g$ in the force regimes examined in this study (Fig. 10A). Passive bundles undergo rapid unsustained elongation even for low values of $f_{\text{ext}}$ (Fig. 10B). The elongated structures do not quickly return to their starting value of $R_g$ once external forces are removed.

#### B. Active structures

Active asters undergo repeated unsustained elongation for force values which are large enough to break the aster's radial symmetry (Fig. 11A). Asters quickly return to their starting configuration once external forces are removed. Active bundles undergo sustained elongation at small values of $f_{\text{ext}}$, and unsustained elongation for larger values of $f_{\text{ext}}$ (Fig. 11B). Upon the removal of forces, bundles return to their initial steady state length.

Asters with cross-linkers with higher activity than those in Fig. 11A also undergo unsustained elongation. Due to the higher velocity of motors, the averaged length of the structure under forces is shorter than for $v_0 = 0.4$ (Fig. 12A). Bundles with cross-linkers with higher activity also undergo elongation (Fig. 12B). Due to the higher velocity of motors, at larger forces the elongated bundles begin undergoing more rapid breakup, which is captured by a decreasing averaged $R_g$ for $f_{\text{ext}} > 0.015$.

### VII. DYNAMIC STEADY STATE OF ACTIVE EXTENSILE BUNDLES

Following the framework of Kruse & Jülicher [4], we describe the bundle in terms of densities $c^+(x,t)$ and $c^-(x,t)$ of plus- and minus-oriented filaments along the bundle axis $x \in [0, L]$. Extending their Eq. 1 to include filament turnover, the equations of motion become:

$$\partial_t c^+ = D\partial_x^2 c^+ - \partial_x J^{++} - \partial_x J^{-+} + k_{\text{in}} c^+_{\text{res}} - k_{\text{out}}\, c^+ \tag{3}$$

$$\partial_t c^- = D\partial_x^2 c^- - \partial_x J^{+-} - \partial_x J^{--} + k_{\text{in}} c^-_{\text{res}} - k_{\text{out}}\, c^- \tag{4}$$

where $J^{\alpha\beta}$ are motor-mediated fluxes between filament pairs of type $(\alpha, \beta)$, $k_{\text{in}}$ is the rate of filament recruitment from the bundle periphery with reservoir density $c^{\pm}_{\text{res}}$, and $k_{\text{out}}$ is the spontaneous filament unbinding rate. We shall assume this spontaneous unbinding to be insignificant since each filament is connected to many neighbouring filaments on an average. The major contribution to filament loss comes from the motor driven fluxes at the bundle ends where motor-driven sliding expels filaments from the overlap zone.

As argued in the main text, parallel pair fluxes ($J^{++}$, $J^{--}$) generate no net sliding. The dominant motor-mediated contribution comes from anti-parallel pairs, with flux:

$$J^{-+} \sim \rho_m v_0\, c^+ c^- \tag{5}$$

where $\rho_m$ is the motor density. Assuming the steady state bundle have a fixed total number of filaments, leads to $\partial_t c^+ + \partial_t c^- = 0$. Now integrating Eq. (3) & (4) over the bundle length $L$:

$$\int_0^L (\partial_x J^{-+} + \partial_x J^{+-})\, dx = \int_0^L k_{\text{in}}(c^+_{\text{res}} + c^-_{\text{res}})\, dx \tag{6}$$

The left-hand side evaluates to the net flux of filaments expelled at the bundle ends by anti-parallel motor sliding:

$$\left[J^{-+} + J^{+-}\right]_0^L = k_{\text{in}}\,(c^+_{\text{res}} + c^-_{\text{res}}) \cdot L \tag{7}$$

Eq. (7) shows the dynamic steady state where the total rate of filament expulsion from the bundle ends due to anti-parallel motor sliding is balanced by the total rate of filament recruitment over the bundle length.

### VIII. SELF-ASSEMBLY MECHANISM AND CROSSLINKER AND MOTOR DENSITY VARIATION

Passive asters ($v_0 = 0$) form through crosslinker-mediated mechanical aggregation. At low crosslinker stiffness ($k = 2$ pN $\mu$m$^{-1}$), crosslinkers can bridge filaments over a wide range of relative orientations without incurring large elastic energy penalties ($E_{\text{cross}} = k\, l_m^2/2 \ll k_B T$ for small extensions), driving radial condensation into apolar asters in which filament barbed ends point randomly inward or outward. At higher crosslinker stiffness, bridging is increasingly restricted to nearly parallel filament pairs, favoring bundle formation instead. Active asters ($v_0 > 0$) form through the same crosslinker-mediated aggregation, with the additional effect of directed motor walking driving polarity sorting — accumulating all barbed ends at the aster center.

The ratio of motors to filaments (1000/200 = 5 motors per filament on average) was chosen to yield well-connected structures with robust steady-state morphologies while remaining computationally tractable. We verified that the key qualitative results — the aster-to-bundle transition and elastic recovery of active structures — are preserved when the number of filaments is varied between 100 and 400, and the motor-to-filament ratio is varied by ±50%.

To further characterize the sensitivity of our results to the number of crosslinkers and motors, we performed additional simulations varying these parameters independently. Increasing the number of passive crosslinkers at fixed filament number drives a morphological transition from no stable structures to asters to bundles (Fig. 18). This confirms that the aster-to-bundle transition is a robust feature of the crosslinker-filament system and not an artifact of our chosen crosslinker number. Decreasing the number of active motors below the baseline value leads to a loss of well-defined aster structures, as the motor density becomes insufficient to maintain the polarity-sorted configuration through directed walking (Fig. 19). These results confirm that the motor number sets a lower bound for the emergence of active cytoskeletal organization, and that the qualitative phenomenology reported in the main text, i.e., aster formation and bundle formation, is robust within the parameter range explored.

| Parameter name | Parameter value | Units |
|---|---|---|
| `time_step` | 0.001 | s |
| `viscosity` | 0.001 | pN s µm$^{-2}$ |
| `temperature` ($k_B T$) | 0.0042 | pN µm |
| simulation box size | $5 \times 5$ | µm$^2$ |
| filament `rigidity` | 0.1 | pN µm$^2$ |
| filament `segmentation` | 0.1 | µm |
| filament `length` | 0.25 | µm |
| number of filaments | 200 | — |
| filament confinement | 1 | pN µm$^{-1}$ |
| myosin `binding_rate` | 10 | s$^{-1}$ |
| myosin `binding_range` | 0.1 | µm |
| myosin `unbinding_rate` | 1 | s$^{-1}$ |
| myosin `unbinding_force` | 10 | pN |
| myosin `activity` | `move` | (towards actin + end) |
| myosin `stall_force` ($f_0$) | 10 | pN |
| myosin `unloaded_speed` ($v_0$) | 0.8, 1.0, 1.2 | µm s$^{-1}$ |
| myosin `stiffness` ($k$) | 0–10 | pN µm$^{-1}$ |
| myosin rest `length` | 0 | µm |
| number of myosin motors | 1000 | — |

TABLE I: Simulation parameters.

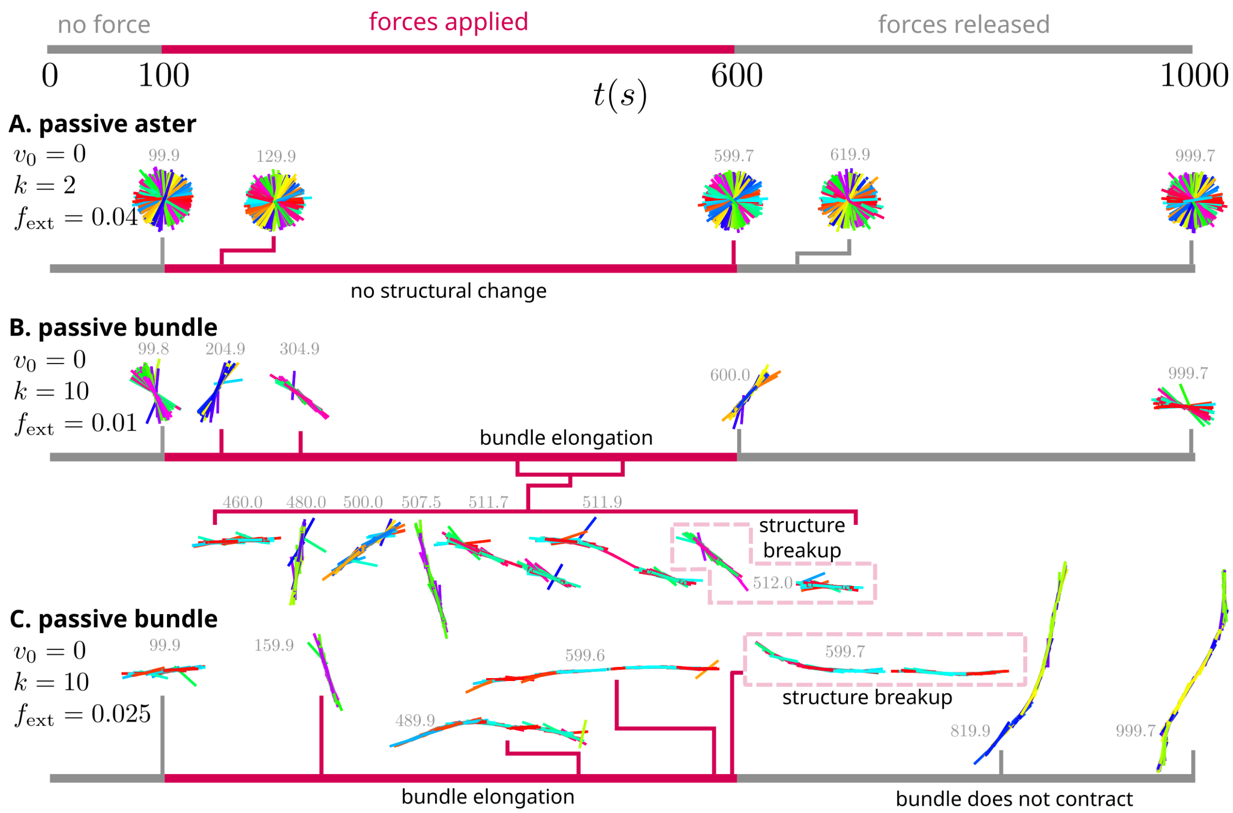


FIG. 1: Graphical summary of simulation results, passive system ($v_0 = 0$). Gray portions of simulation timelines correspond to regimes with no applied external forces. Red portions of timelines correspond to applied forces. Exact timestamps are shown in gray next to each snapshot. (A) Passive asters do not respond to the external forces in the regime that we examined. (B–C) Passive bundles elongate upon the application of external force, but do not contract upon the removal of forces.

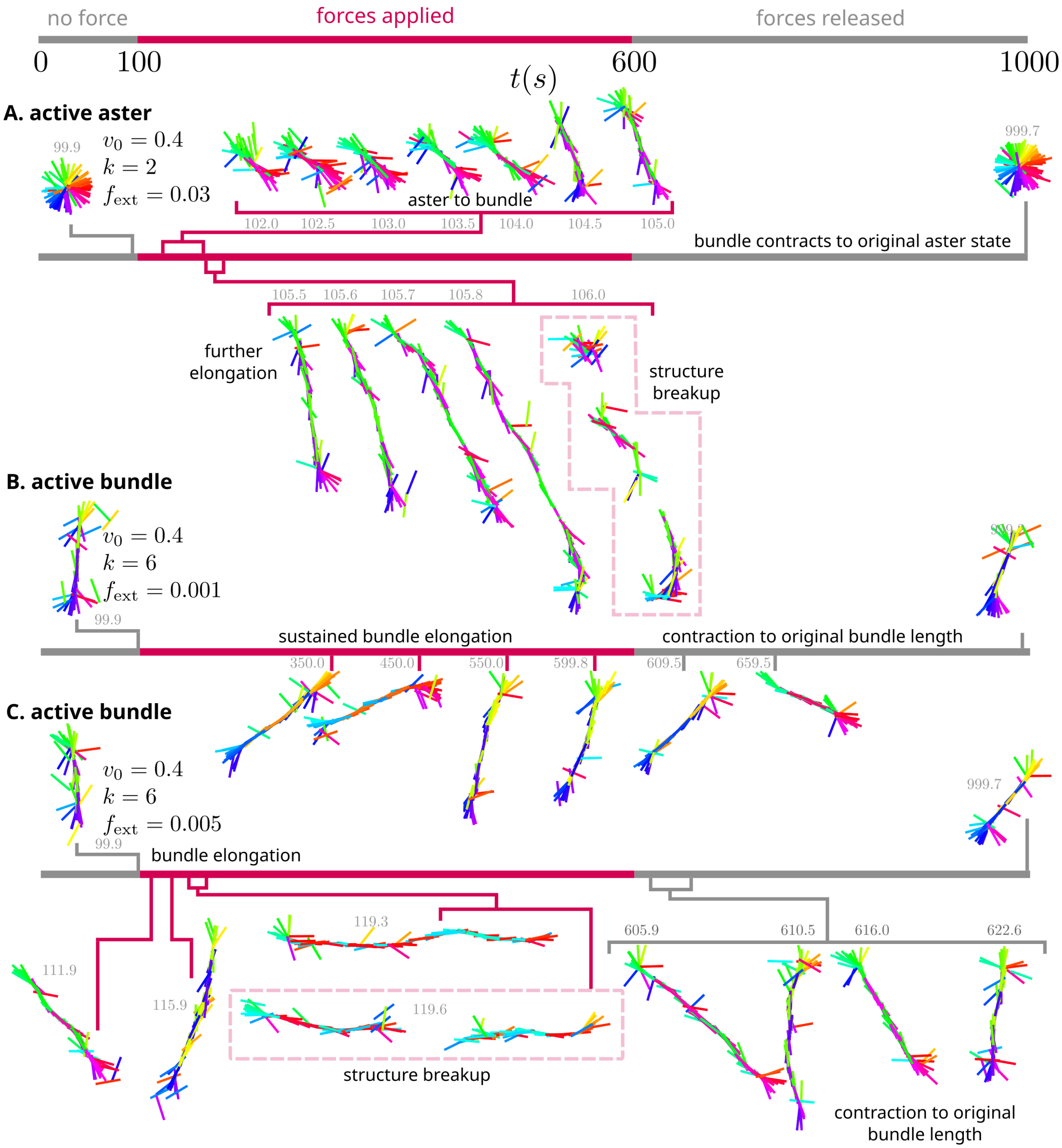


FIG. 2: Graphical summary of simulation results, active system ($v_0 = 0.4$). Gray portions of simulation timelines correspond to regimes with no applied external forces. Red portions of timelines correspond to applied forces. Exact timestamps are shown in gray next to each snapshot. (A) Active asters elongate into bundles for larger $f_{ext}$ but eventually break up. (B) Active bundles undergo sustained elongation for smaller values of $f_{ext}$. (C) Active bundles undergo unsustained elongation for larger $f_{ext}$.

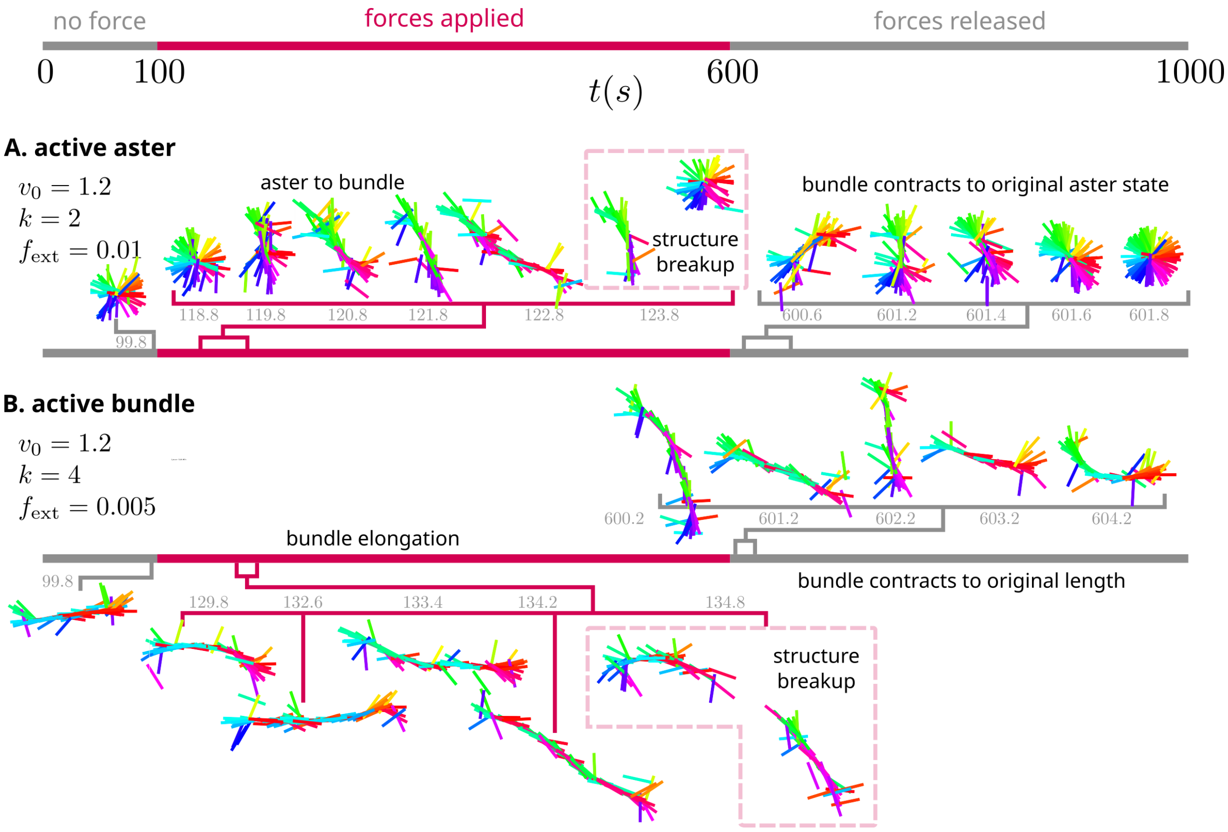


FIG. 3: Graphical summary of simulation results, active system ($v_0 = 1.2$). Gray portions of simulation timelines correspond to regimes with no applied external forces. Red portions of timelines correspond to applied forces. Exact timestamps are shown in gray next to each snapshot. (A) Active structures initially elongate and then break up. (B) Active bundles can undergo sustained elongation at low force values.

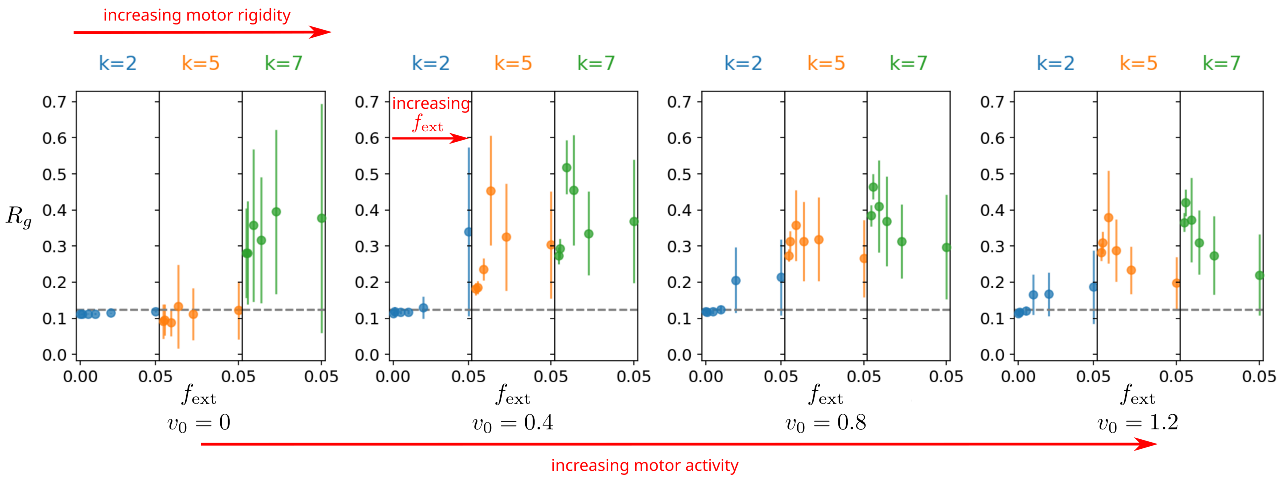


FIG. 4: Radius of gyration of the largest cluster under varying activity conditions.

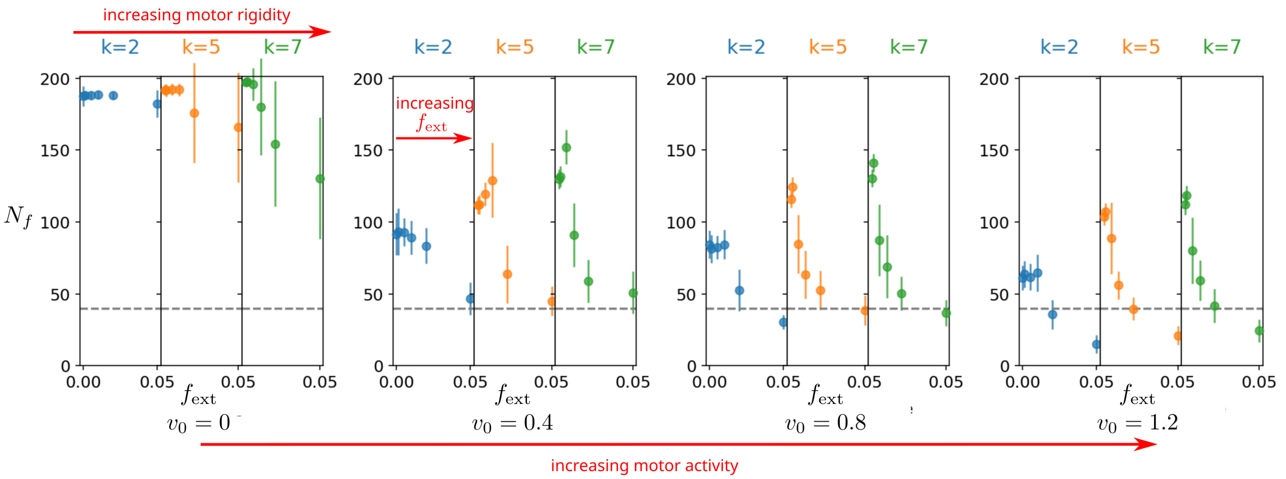


FIG. 5: Number of filaments in the largest cluster under varying activity conditions.

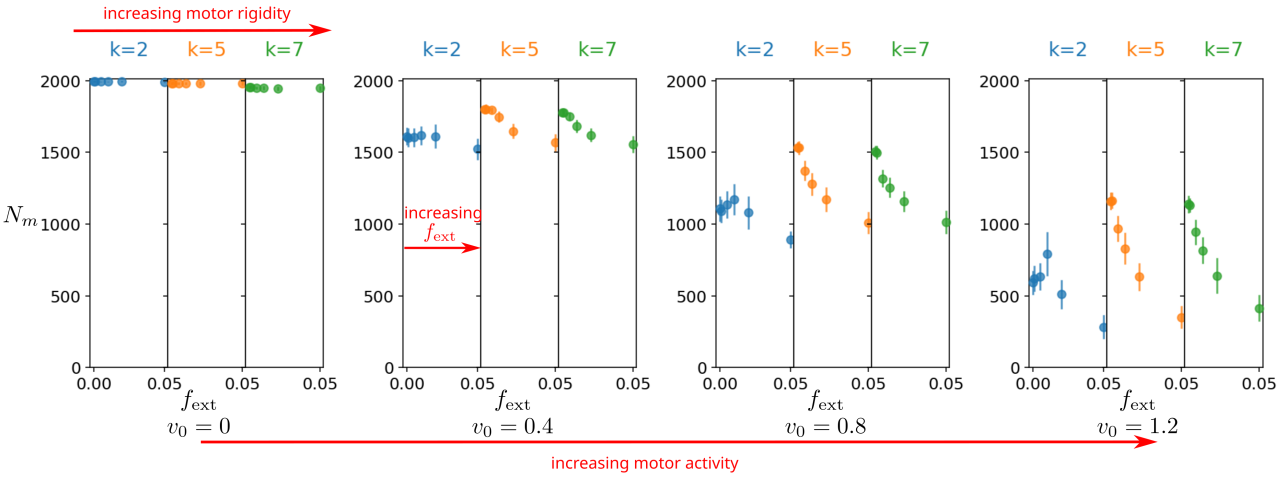


FIG. 6: Average number of motors in the largest cluster under varying activity conditions.

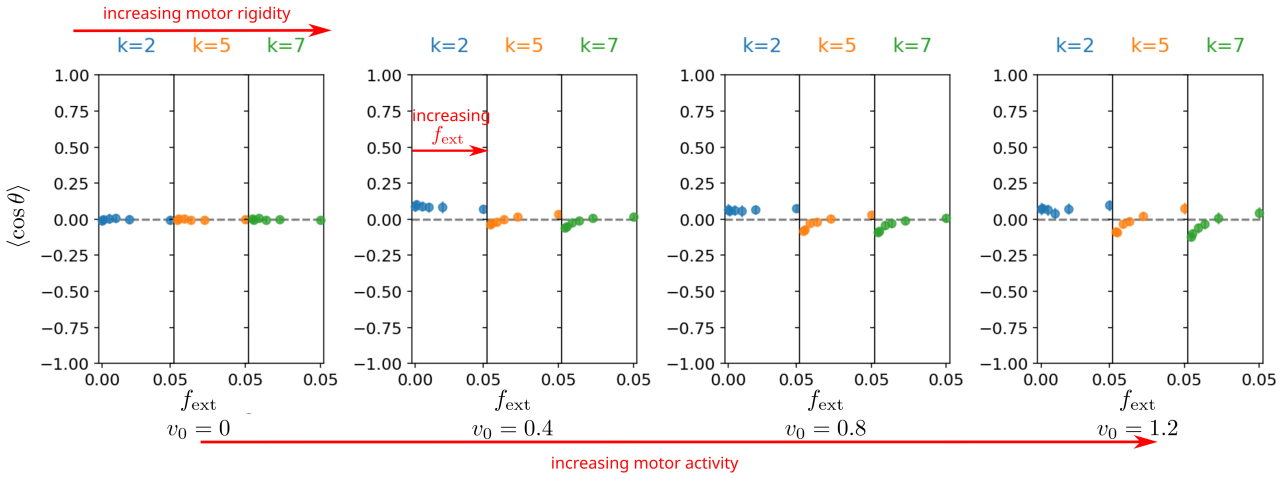


FIG. 7: $\langle \cos\theta \rangle$: average filament-motor angles in the largest cluster under varying activity conditions.

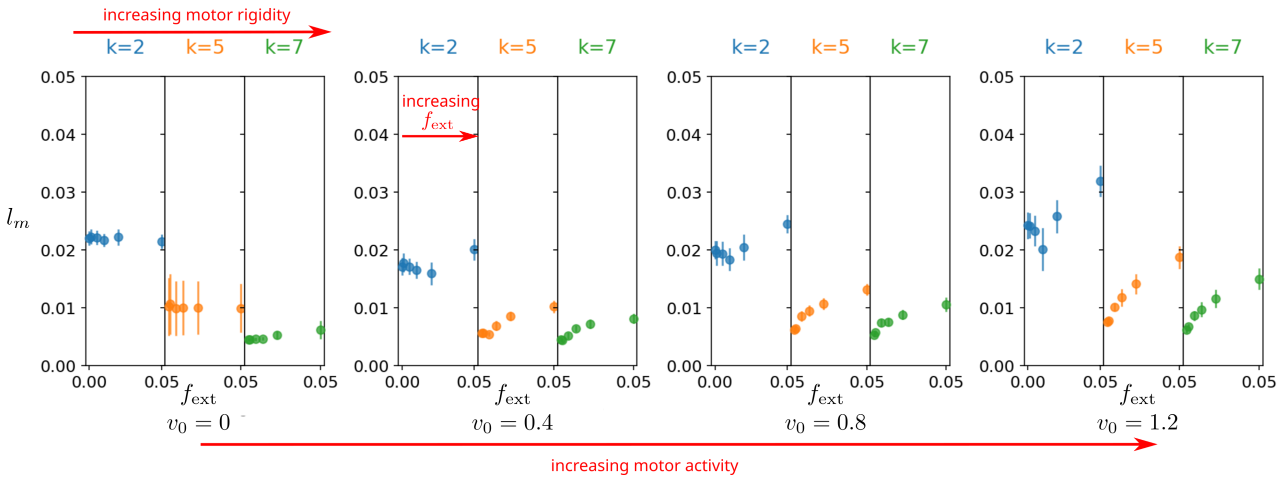


FIG. 8: Average motor length in the largest cluster under varying activity conditions.

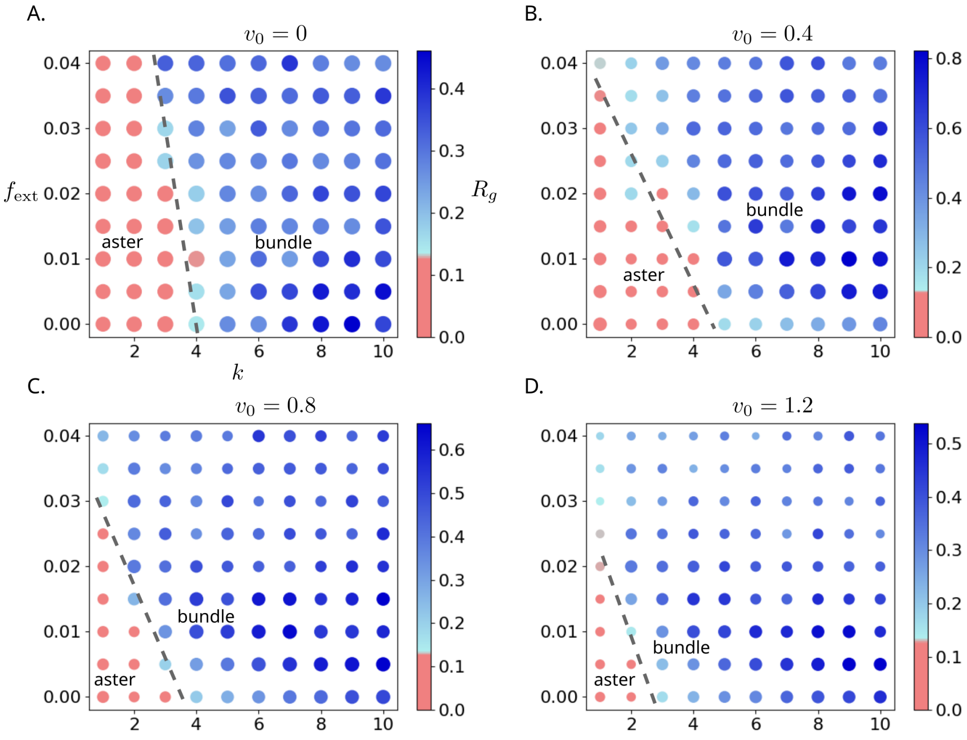


FIG. 9: Response of structures to external forces depends on the activity level of the cross-linking motors. Colorbar corresponds to the radius of gyration $R_g$ of largest structure in the system. Point sizes are proportional to the number of filaments in the largest structure. (A) $v_0 = 0$, passive; (B) $v_0 = 0.4$, active; (C) $v_0 = 0.8$, active; (D) $v_0 = 1.2$, active.

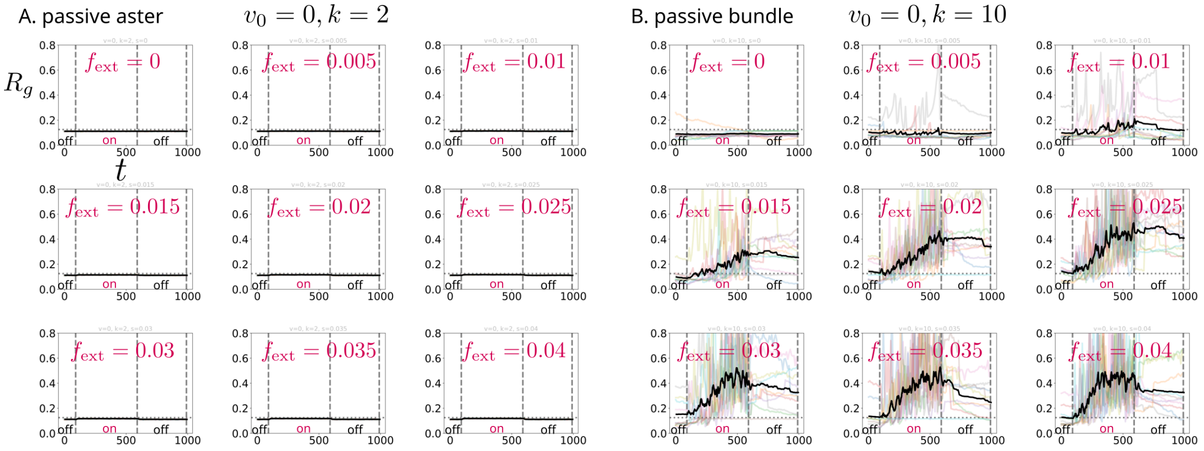


FIG. 10: Passive, $v_0 = 0$. Traces of $R_g$ for structures under various external forces. Color lines represent traces for individual trajectories ($N = 10$), and heavy black line indicates a moving average across all trajectories in the ensemble ($N = 50$, averaging time 0.1 s). Vertical dashed lines indicate boundaries between force regimes. Horizontal dashed line indicates $R_g = 0.125$, corresponding to the radius of gyration of a "perfect" aster.

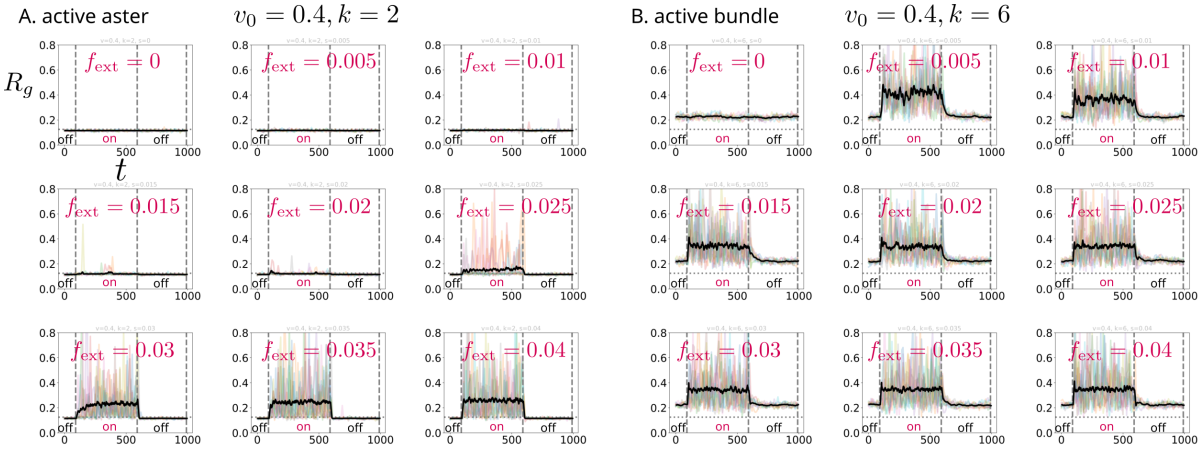


FIG. 11: Active, $v_0 = 0.4$. Traces of $R_g$ for structures under various external forces. Color lines represent traces for individual trajectories ($N = 10$), and heavy black line indicates a moving average across all trajectories in the ensemble ($N = 50$, averaging time 0.1 s). Vertical dashed lines indicate boundaries between force regimes. Horizontal dashed line indicates $R_g = 0.125$, corresponding to the radius of gyration of a "perfect" aster.

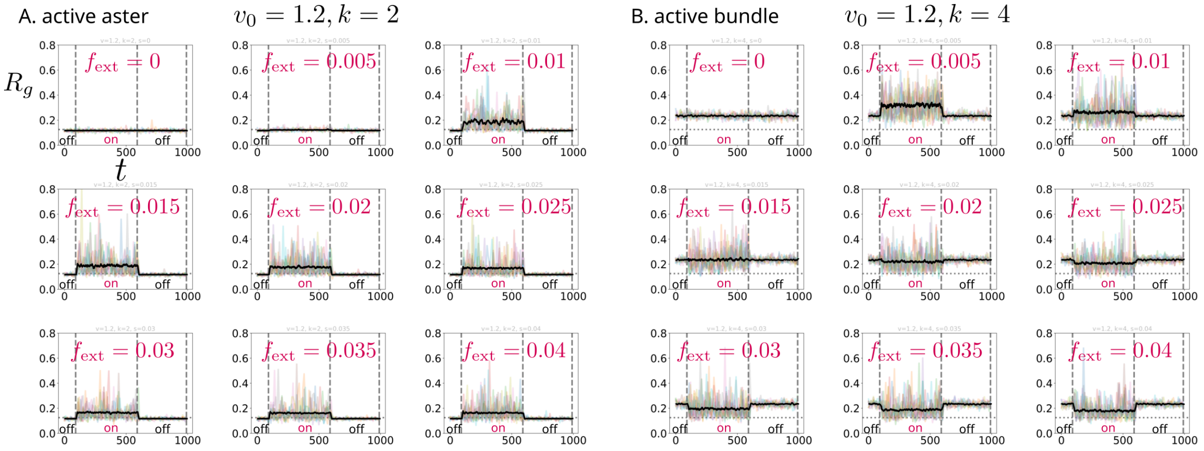


FIG. 12: Active, $v_0 = 1.2$. Traces of $R_g$ for structures under various external forces ("off" labels indicate no applied $f_{\rm ext}$, red "on" labels indicate applied $f_{\rm ext}$). Color lines represent traces for individual trajectories ($N = 10$), and heavy black line indicates a moving average across all trajectories in the ensemble ($N = 50$, averaging time 0.1 s). Vertical dashed lines indicate boundaries between force regimes. Horizontal dashed line indicates $R_g = 0.125$, corrisponding to the radius of gyration of an "ideal" aster.

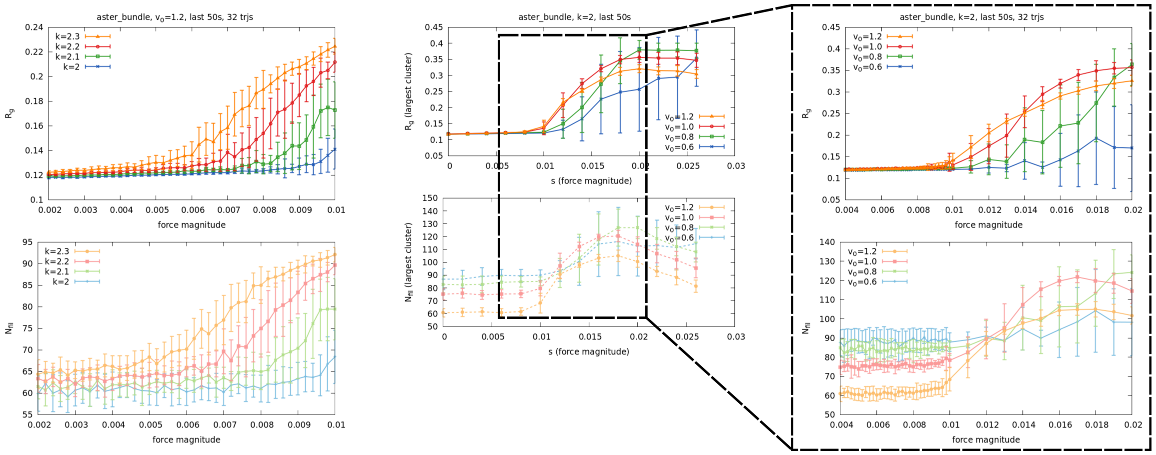


FIG. 13: **Aster threshold forces.** Variation of minimum threshold force required to pull an aster into a bundle for various motor properties (left: motor rigidity $k$; center and right: unloaded motor velocity $v_0$).

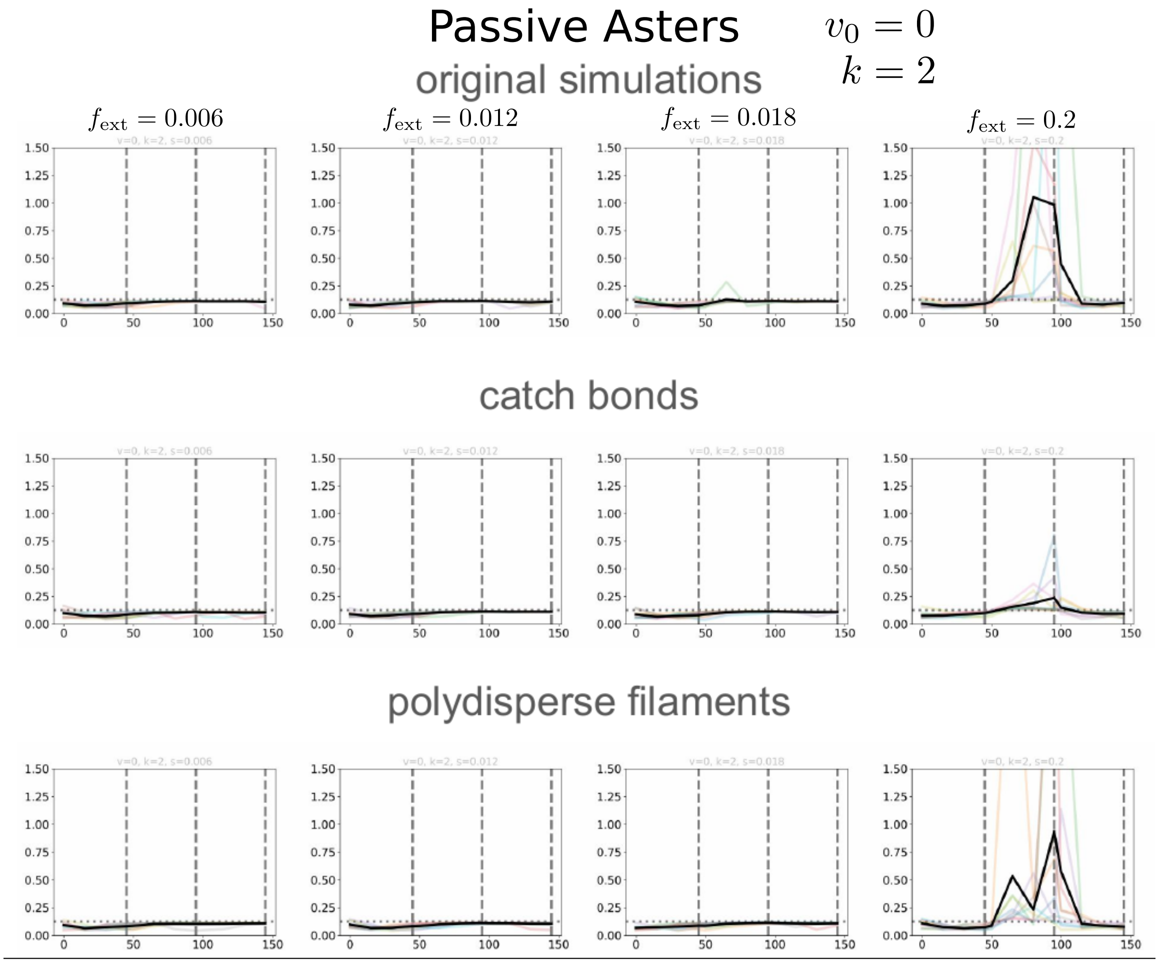


FIG. 14: **Effect of catch bond in crosslinker unbinding and polydispersity of filament size on passive asters.** (top) Original simulations with slip bond behaviour and monodisperse filaments. (middle) Force-dependent morphological change in passive asters for crosslinkers with catch bond behaviour. (bottom) Force-dependent morphological change in passive asters with polydispersity in filament size.

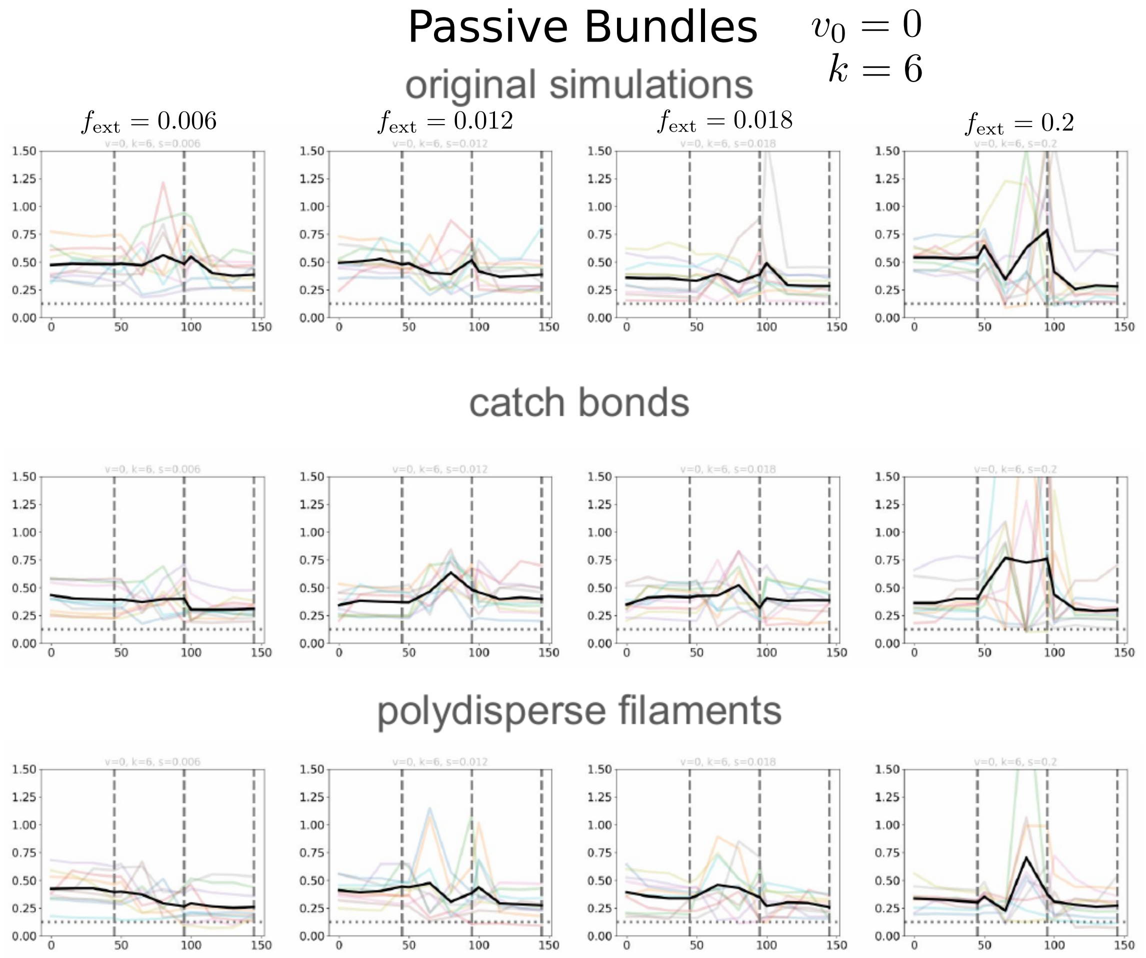


FIG. 15: **Effect of catch bond in crosslinker unbinding and polydispersity of filament size on passive bundle.** (top) Original simulations with slip bond behaviour and monodisperse filaments. (middle) Force-dependent morphological change in passive bundles for crosslinkers with catch bond behaviour. (bottom) Force-dependent morphological change in passive bundles with polydispersity in filament size.

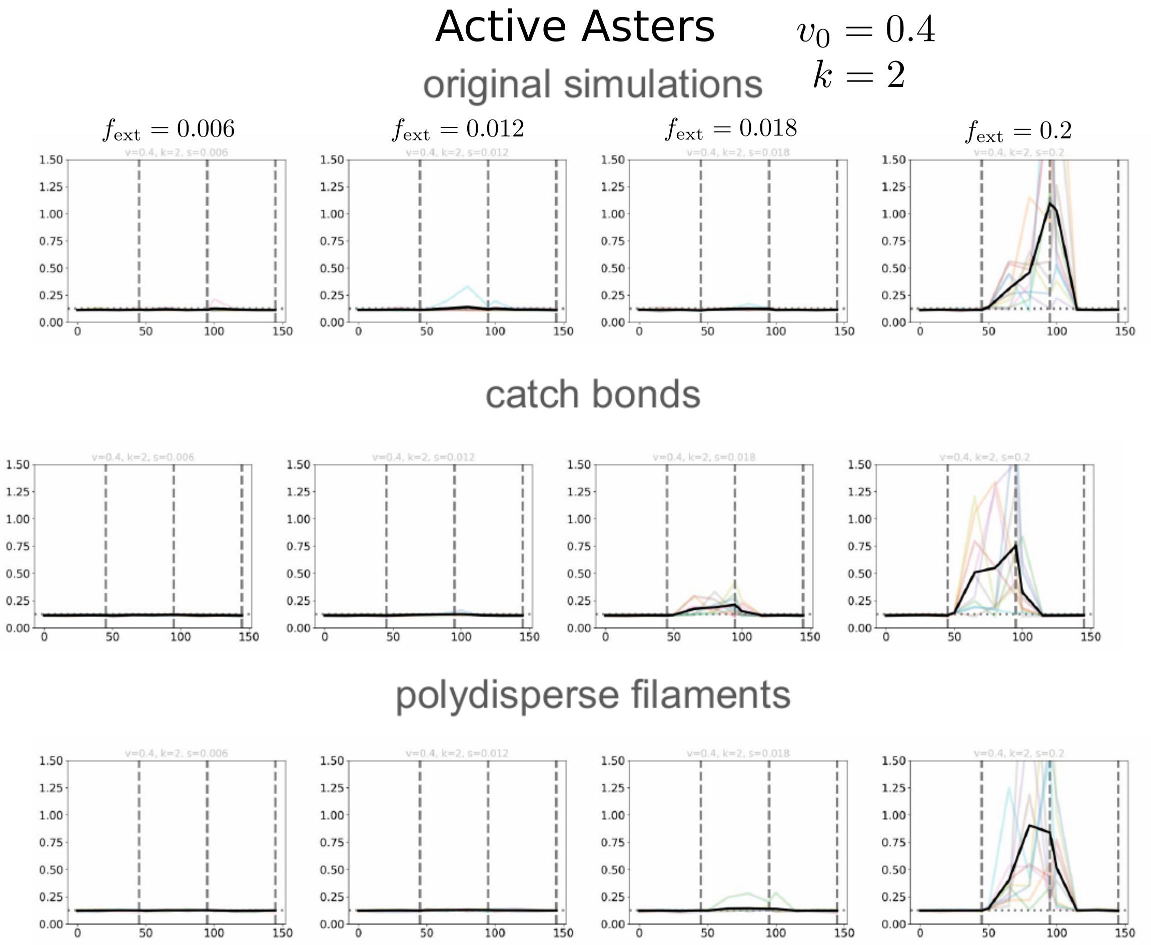


FIG. 16: **Effect of catch bond in crosslinker unbinding and polydispersity of filament size on active asters.** (top) Original simulations with slip bond behaviour and monodisperse filaments. (middle) Force-dependent morphological change in active asters for crosslinkers with catch bond behaviour. (bottom) Force-dependent morphological change in active asters with polydispersity in filament size.

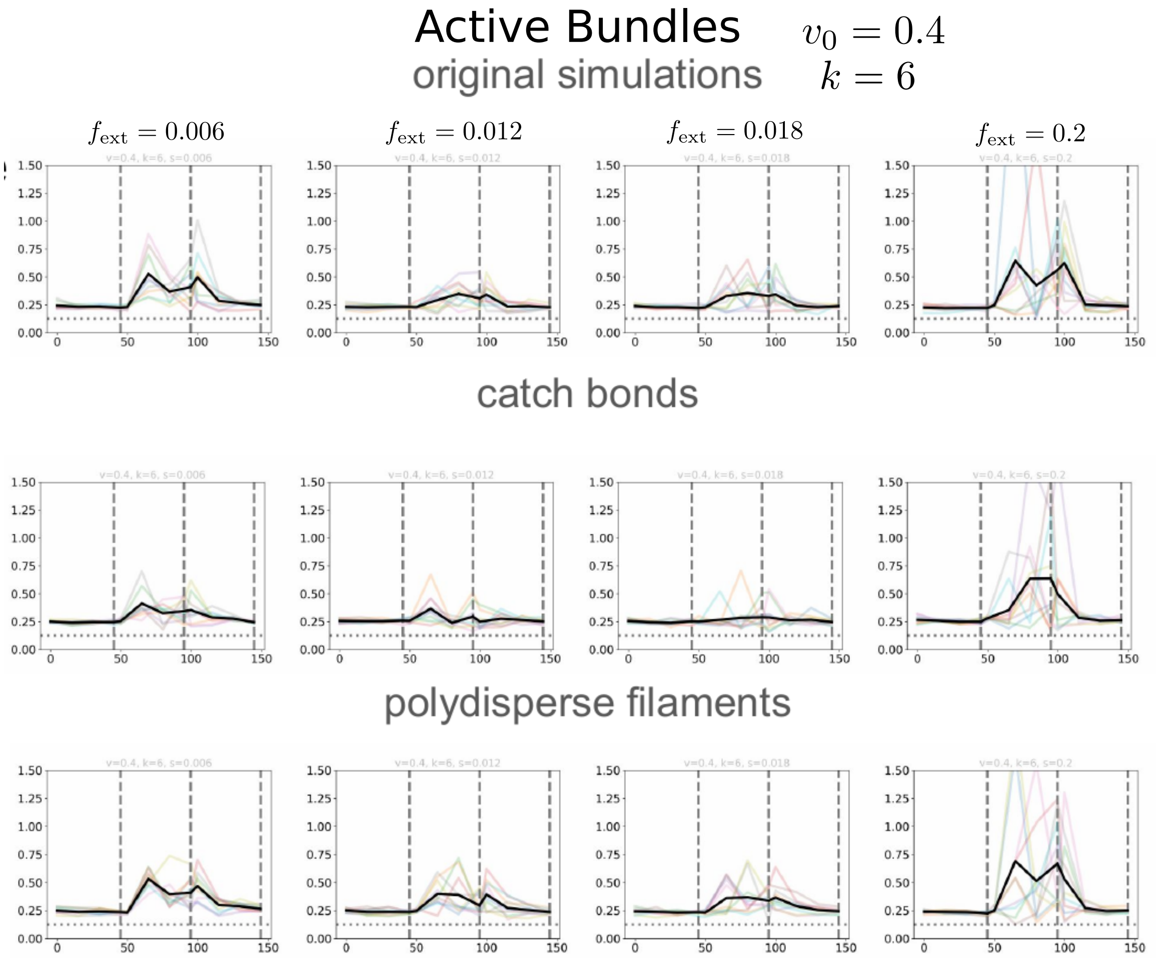


FIG. 17: **Effect of catch bond in crosslinker unbinding and polydispersity of filament size on active bundle.** (top) Original simulations with slip bond behaviour and monodisperse filaments. (middle) Force-dependent morphological change in active bundles for crosslinkers with catch bond behaviour. (bottom) Force-dependent morphological change in active bundles with polydispersity in filament size.

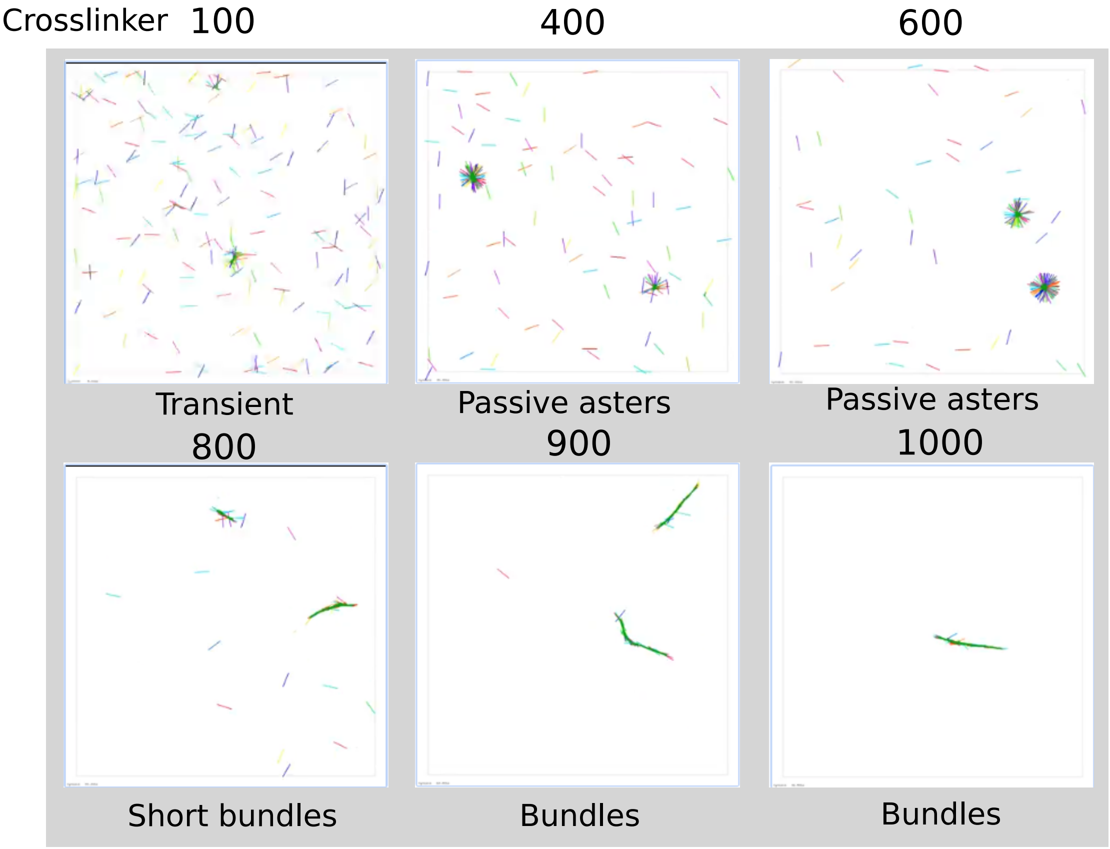


FIG. 18: **Effect of changing crosslinker density on passive aster.** Increasing crosslinker number shows transitions from no stable structure to aster and bundle in very high crosslinker density.

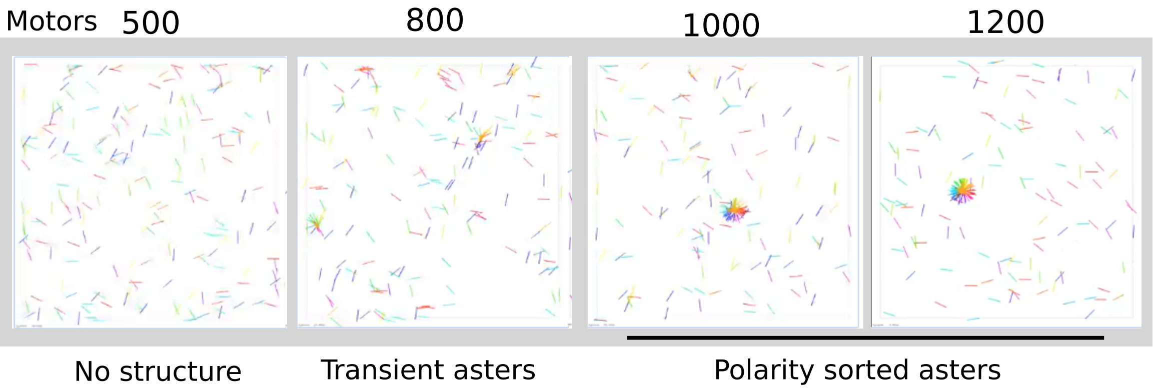


FIG. 19: **Effect of changing motor density on active aster.** Increasing motor number shows transitions from no stable structure to transient and stable asters in high enough motor density.

[1] Francois Nedelec and Dietrich Foethke. Collective langevin dynamics of flexible cytoskeletal fibers. *New Journal of Physics*, 9(11):427–427, Nov 2007.
[2] ctypes — A foreign function library for Python — docs.python.org. https://docs.python.org/3/library/ctypes.html.
[3] Python interface to cytosim for introducing external forces. https://gitlab.com/alamda/cytosim/-/tree/python_forces.
[4] Karsten Kruse and F Jülicher. Actively contracting bundles of polar filaments. *Physical review letters*, 85(8):1778, 2000.
